# Redox response of iron-sulfur glutaredoxin GRXS17 activates its holdase activity to protect plants from heat stress

**DOI:** 10.1101/2020.01.07.896506

**Authors:** Laura Martins, Johannes Knuesting, Laetitia Bariat, Avilien Dard, Sven A. Freibert, Christophe H. Marchand, David Young, Nguyen Ho Thuy Dung, Anne Debures, Julio Saez-Vasquez, Stéphane D. Lemaire, Roland Lill, Joris Messens, Renate Scheibe, Jean-Philippe Reichheld, Christophe Riondet

**Author notes:** To whom correspondence should be addressed. Jean-Philippe Reichheld. Laboratoire Génome et Développement des Plantes, Université Perpignan Via Domitia, F-66860 Perpignan, France. These authors contributed equally to this work (Laura Martins, Johannes Knuesting).

## Abstract

Living organisms use a large panel of mechanisms to protect themselves from environmental stress. Particularly, heat stress induces misfolding and aggregation of proteins which are guarded by chaperone systems. Here, we examine the function the glutaredoxin GRXS17, a member of thiol reductases families in the model plant *Arabidopsis thaliana*. GRXS17 is a nucleocytosolic monothiol glutaredoxin consisting of an N-terminal thioredoxin (TRX)-domain and three CGFS-active site motif-containing GRX-domains that coordinate three iron-sulfur (Fe-S) clusters in a glutathione (GSH)-dependent manner. As a Fe-S cluster-charged holoenzyme, GRXS17 is likely involved in the maturation of cytosolic and nuclear Fe-S proteins. In addition to its role in cluster biogenesis, we showed that GRXS17 presents both foldase and redox-dependent holdase activities. Oxidative stress in combination with heat stress induces loss of its Fe-S clusters followed by subsequent formation of disulfide bonds between conserved active site cysteines in the corresponding TRX domains. This oxidation leads to a shift of GRXS17 to a high-MW complex and thus, activates its holdase activity. Moreover, we demonstrate that GRXS17 is specifically involved in plant tolerance to moderate high temperature and protects root meristematic cells from heat-induced cell death. Finally, we showed that upon heat stress, GRXS17 changes its client proteins, possibly to protect them from heat injuries. Therefore, we propose that the iron-sulfur cluster enzyme glutaredoxin GRXS17 is an essential guard to protect proteins against moderate heat stress, likely through a redox-dependent chaperone activity. All in all, we reveal the mechanism of an Fe-S cluster-dependent activity shift, turning the holoenzyme GRXS17 into a holdase that prevents damage caused by heat stress.

## INTRODUCTION

Glutaredoxins (GRX) are small ubiquitous thiol reductases belonging to the thioredoxin (TRX) superfamily (Rouhier et al., 2008; Meyer et al., 2009). Their thiol-disulfide exchange capacity generally depends on glutathione as a reductant, and relies on an active site harboring at least one cysteine. In plants, glutaredoxins are organized in multigene families including four different subgroups (Meyer et al., 2012). In Arabidopsis, members of the type-I and -II subfamilies are characterized by their capacity to bind a labile iron-sulfur cluster (Fe-S cluster), coordinated by conserved active site amino acids (CG/SF/YS/C) and a glutathione molecule, a feature largely conserved within eukaryotic organisms (Lillig et al., 2005; Wingert et al., 2005; Rouhier et al., 2007; Bandyopadhyay et al., 2008; Couturier et al., 2011; Riondet et al., 2012; Berndt and Lillig, 2017). Remarkably, all type-II GRX exhibit a CGFS active site and most of them are able to transfer their Fe-S cluster to apo-enzymes *in vitro*, indicating a role of these GRX as Fe-S cluster donors for Fe-S-containing enzymes (Bandyopadhyay et al., 2008; Moseler et al., 2015). Consistent with this activity, most plant type-II GRX are able to rescue defects of the yeast *grx5* mutant deficient in mitochondrial Fe-S cluster biogenesis (Bandyopadhyay et al., 2008; Moseler et al., 2015; Knuesting et al., 2015). The labile character of the Fe-S cluster in type-I and -II GRX suggests that the holo- and apo-forms might perform different activities. In absence of the Fe-S cluster, some type-I GRXs reduce disulfides in the catalytic cycle of enzymes, such as in methionine sulfoxide reductase and peroxiredoxins, or the mixed-disulfide between glutathione and glyceraldehyde-3-phosphate dehydrogenase (GAPDH) (Zaffagnini et al., 2008; Couturier et al., 2011; Riondet et al., 2012). PICOT-like GRX are multi-domain type-II GRX found in the cytosol of most eukaryotic organisms (Berndt and Lillig, 2017). While in most organisms, PICOT proteins are composed of one or two adjacent GRX domains with a CGFS active-site sequence motif preceded by a TRX domain, the PICOT proteins of higher plants, like GRXS17, contain a third CGFS-containing GRX domain (Couturier et al., 2009). *In vitro*, GRXS17 is a homodimer and each GRX-CGFS domain coordinates an Fe-S cluster in a glutathione(GSH)-dependent manner (Knuesting et al., 2015). GRXS17 is capable of complementing the defect of the yeast *grx5* deletion mutant by transferring Fe-S clusters to mitochondrial acceptor proteins (Bandyopadhyay et al., 2008; Knuesting et al., 2015). Affinity chromatography approaches with crude extracts have shown that GRXS17 associates with known cytosolic iron-sulfur protein assembly (CIA) components like Met18, Dre2, and with other cytosolic Fe-S cluster-containing proteins like xanthine dehydrogenase 1 (XDH1) and thiouridylase subunits 1 and 2 (Iñigo et al., 2016; Wang et al., 2019). GRXS17 can be additionally found in the nucleus, and it interacts with cytosolic BolA2 (Couturier et al., 2014) and with NF-YC11, a subunit of the NF-Y transcription factor complex (Knuesting et al., 2015). However, for most of its partners, the role of the Fe-S cluster remained enigmatic. Genetic evidence from different plant species suggests a protective role of GRXS17 during thermal, genotoxic and drought stress (Cheng et al., 2011; Wu et al., 2012; Wu et al., 2017; Iñigo et al., 2016; Hu et al., 2017), in meristem development (Knuesting et al., 2015), hormonal responses (Cheng et al., 2011), and during redox-dependent processes (Yu et al., 2018).

Here, we focus on the redox-dependent properties of GRXS17 and its role in the cell. When subjected to oxidative conditions, reconstituted holo-GRXS17 rapidly loses its Fe-S cluster and becomes a cysteine-dependent holdase. The switch to holdase involves oligomerization. Moreover, GRXS17 exhibits foldase activity which does not depend on its active-site cysteines. To confirm the *in vivo* importance of these chaperone activities of GRXS17, we analysed a *grxs17* knock-out mutant that was highly sensitive to heat stress. Mutant complementation experiments with several cysteine variants of GRXS17 indicate the crucial role of the active-site cysteines in protecting proteins during heat stress. Finally, GRXS17 associates with a different set of partner proteins after heat stress, indicating a stress-dependent functional switch of GRXS17 that could contribute to protecting cells from heat stress damage.

## RESULTS

### GRXS17 releases its Fe-S cluster and oligomerizes under combined oxidative and heat stress

When purified under anaerobic conditions, GRXS17 adopts a dimeric form coordinating up to three Fe-S clusters together with six molecules of GSH (Knuesting et al., 2015). As Fe-S clusters are highly sensitive to oxidation, we sought to examine the structural modifications of GRXS17 upon transfer to aerobic conditions. As expected, anaerobically reconstituted GRXS17 Fe-S cluster shows a circular dichroism (CD) spectrum typical for the presence of [2Fe-2S] clusters (Figure 1A). When exposed to air, the spectral footprint of the Fe-S cluster was progressively destabilized by oxygen and finally lost, confirming the sensitivity of the Fe-S cluster for oxidation (Figure 1A).

**Figure 1:**
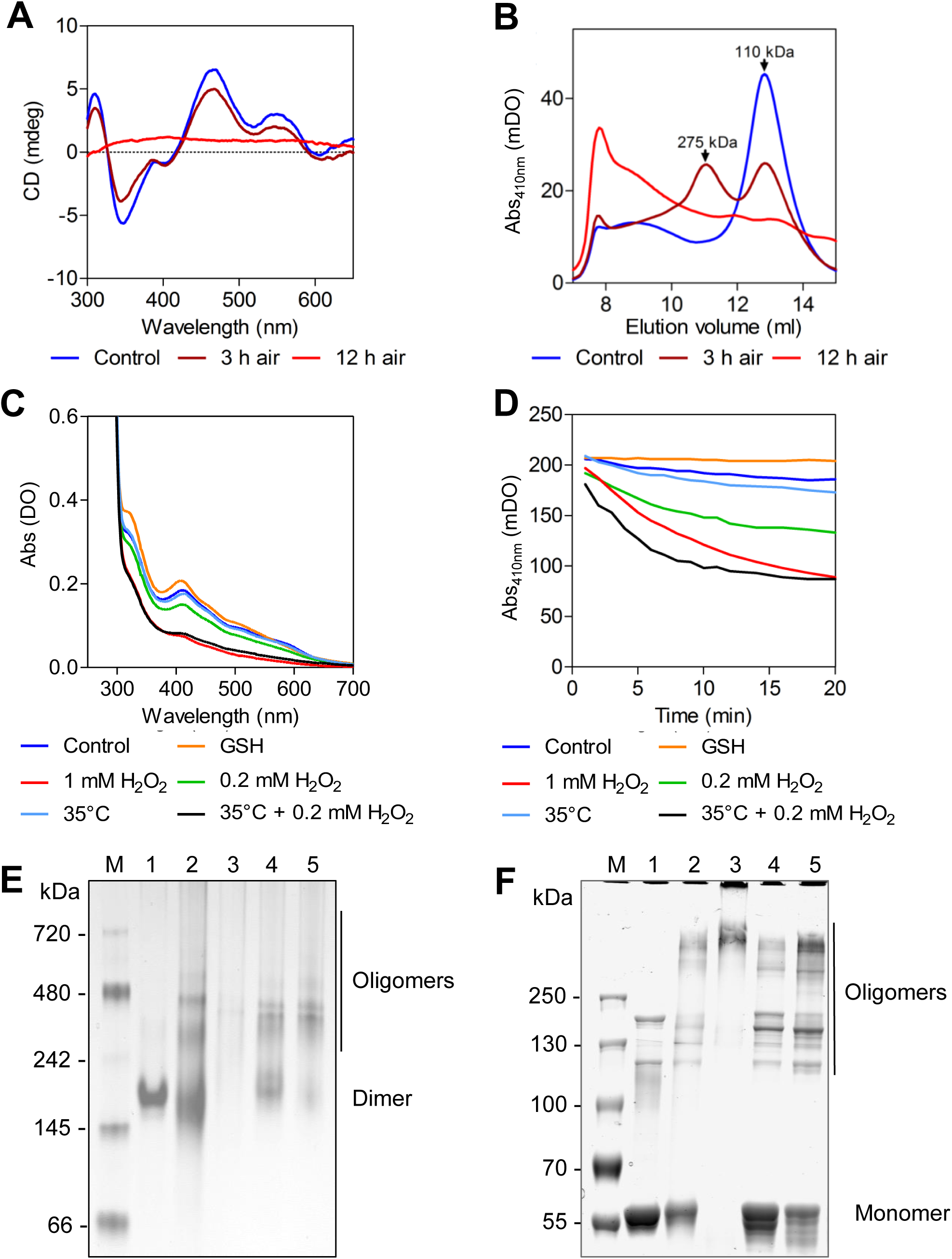
Fe-S cluster stability and oligomerization of reconstituted GRXS17. **(A)** Visible Circular Dichroism (CD)-spectra of reconstituted holo-GRXS17 under reducing conditions (blue) and after oxygen treatment (3h brown; 12h red) at room temperature. **(B)** Size-exclusion chromatography (Sephacryl S300 HR) under anaerobic conditions with reconstituted GRXS17 and after the respective oxygen treatments. **(C)** Absorption spectra of GRXS17 subjected to H_2_O_2_ and heat treatments. UV-VIS absorption spectra were measured after *in vitro* reconstitution in under anaerobic conditions and after 20 min at 20°C (blue), GSH (2.5 mM orange), H_2_O_2_ treatment at 20°C (0.2 mM green; 1 mM red), heat treatment at 35°C in presence of 0.2 mM H_2_O_2_ (black) or without H_2_O_2_ (light blue). **(D)** Absorption at 410 nm of GRXS17 subjected to the same treatments as in **(C)**. **(E-F)** Migration of 5 µg of recombinant GRXS17 with or without the cluster in (**E**) native PAGE or (**F**) in non-reducing SDS-PAGE gels. M, Molecular marker. HoloGRXS17 (1) was maintained on air for 3h (2) or 12h (3), of treated for 10 min at 35°C (4) or with 0.2 mM H_2_O_2_ (5). Samples were subjected to non-reducing SDS-PAGE and protein bands were stained by Coomassie blue. Each experiment is representative of three biological repetitions.

In parallel to the loss of the Fe-S cluster, from size-exclusion chromatography (SEC) assay, we observed a progressive shift of GRXS17 to a high-MW form of >250 kDa after 3 h of exposure to air. After 12 h, a high-MW complex of a higher range was observed (Figure 1B). Similar data were observed by native gel electrophoresis and non-reducing SDS-PAGE (Figure 1E-F). As expected, holo-GRXS17 migrates, on non-reducing native gel, as a single band likely corresponding to a dimeric form. However, after storage under aerobic conditions for more than 12 h, oligomers were formed, underscoring the high sensitivity of holo-GRXS17 to oxygen (Figure 1E-F). Interestingly, the size of the HMW-complexes was somehow lower under non-reducing SDS-PAGE conditions than on a native gel, suggesting that both weak-bond and disulfide-bond interactions are formed under oligomerization. All in all, we showed that the presence of the Fe-S cluster prevents the oligomerization of GRXS17.

To further examine the stability of the Fe-S cluster under oxidative conditions, we exposed the holoGRXS17 to physiological concentrations (0.2-1 mM) of H_2_O_2_. As previously shown (Knuesting et al., 2015), the UV-VIS spectrum of the reconstituted GRXS17 exhibited absorbance peaks at 320 and 410 nm, characteristic for [2Fe-2S] clusters. Upon H_2_O_2_ treatment, the Fe-S cluster is lost in a dose-dependent manner (∼50% loss in 12 and 5 min at 0.2 mM and 1 mM, respectively), confirming the sensitivity of the Fe-S cluster to oxidation (Figure 1C-D). As the high temperature was previously shown to destabilize Fe-S clusters in other proteins (Yan et al., 2013), we also monitor its stability at high temperatures. Both 35°C and 43°C temperatures had little effect on the stability of the cluster after short-term treatments, indicating that the Fe-S cluster is not heat sensitive (Figure 1C-D and Supplemental Figure 1). However, when combining H_2_O_2_ treatment with physiological high-temperatures, an accelerated loss of the Fe-S cluster (∼50% loss in 3 min at 0.2 mM H_2_O_2_) was observed, showing that higher temperatures potentiate the oxidative effect on Fe-S cluster stability. As expected, extensive oligomerization of GRXS17 was observed by native gel and non-reducing SDS-PAGE gels after H_2_O_2_ treatment, while high temperature exposure had less impact on oligomerization of GRXS17 (Figure 2E-F).

**Figure 2:**
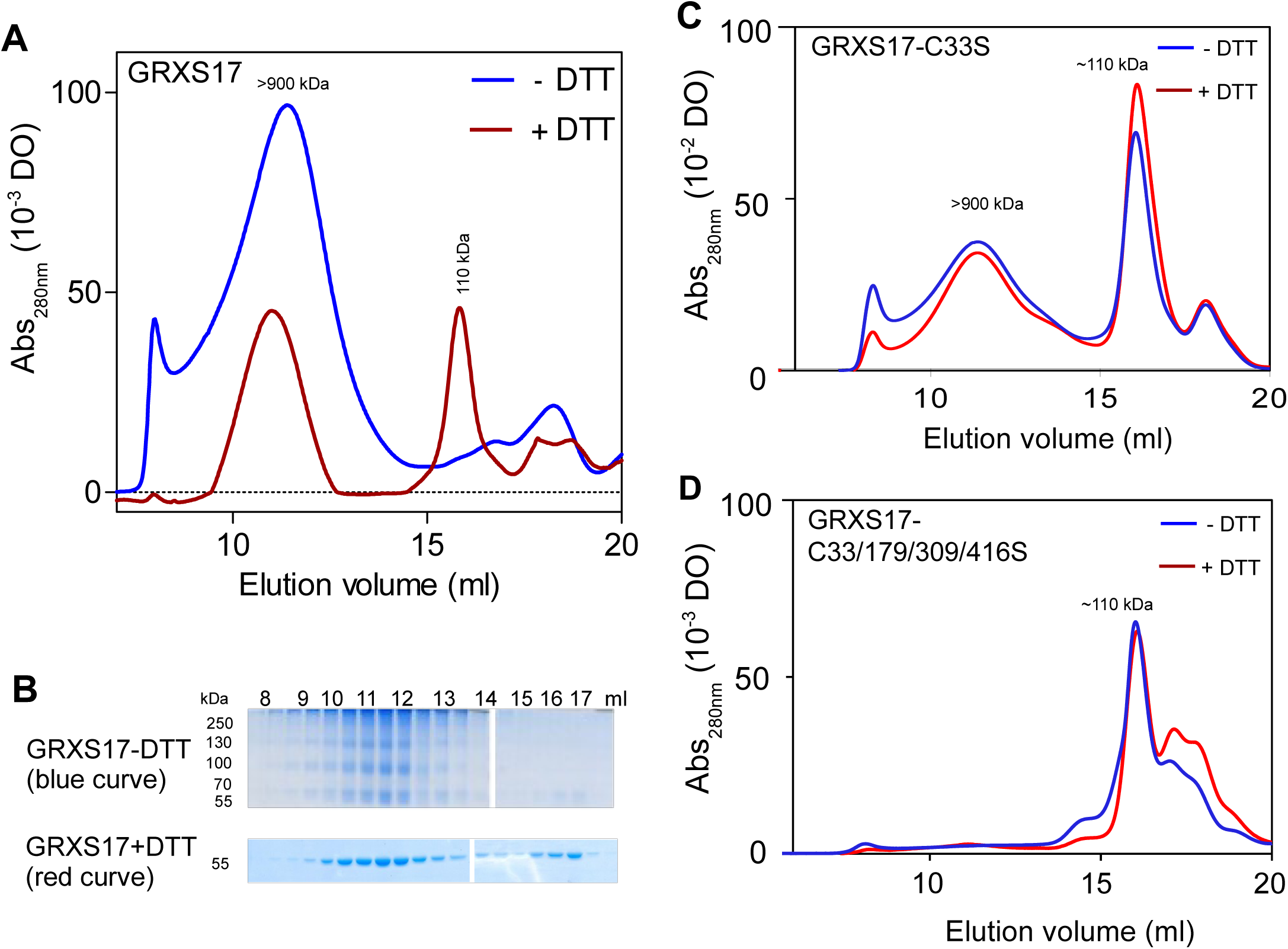
GRXS17 forms high molecular weight complexes. **(A)** Size-exclusion chromatography (SEC) analysis (Superose 6) of GRXS17 protein (740 µg) maintained in air after purification. The protein was reduced with 10 mM DTT (red) or not (blue) before loading on the column. **(B)** Aliquots of elution fraction (30 µL) shown in (A) were subjected to non-reducing SDS-PAGE. **(C-D)** Recombinant GRXS17 mutated in active-site cysteines were maintained in air and subjected to SEC (Superose 6) in presence (red) or absence of 10 mM DTT (blue). **(C)** GRXS17-C33S, **(D)** GRXS17-C33/179/309/416S. Approximative molecular weights are calculated based on the elution volumn of reference proteins (see Methods).

### GRXS17 oligomerizes through intermolecular disulfides and noncovalent interactions

We further explored GRXS17 oligomerization after loss of Fe-S clusters using SEC (Figure 2). As expected, HMW complexes (∼900-1000 kDa) were observed in the native apoGRXS17 (Figure 2A, blue line). When separating these fractions on a non-reducing SDS-PAGE gel, a laddering profile was observed, suggesting that the HMW complexes are partially dissociated by SDS and that the remaining HMW bands contain disulfide-linked GRXS17 oligomers (Figure 2B). These data indicate that native GRXS17 oligomerizes through non-covalent interactions and intermolecular disulfides. Consistently, incubation of the native GRXS17 with DTT prior to SEC led to a decrease of the ∼900-1000 kDa peak and a concomitant increase of the 110 kDa peak, likely corresponding to homodimeric GRXS17 (Figure 2A). Furthermore, SDS-PAGE analyses indicate that the peaks corresponding to oligomeric and dimeric GRXS17 consist of non-covalently interacting monomers (Figure 2B).

In order to analyze the role of the active-site cysteines during the oligomerization, mutated forms of GRXS17 were analyzed by SEC. Interestingly, the TRX-domain C33S mutant behaved similarly to reduced wild-type GRXS17 (Figure 2C), namely exhibiting comparable peaks at ∼900-1000 kDa and 110 kDa. Moreover, the distribution between these peaks was not influenced by DTT treatment prior to SEC, indicating that Cys33 plays a major role in the oligomerization, very likely by forming intermolecular disulfide bonds (Figure 2C). Strikingly, mutation of all active site cysteines completely abolishes oligomerization, as GRXS17-C33/179/309/416S exclusively elutes as a dimer (Figure 2D).

Finally, to verify the impact of Cys mutations on the GRXS17 structure, we analyzed secondary structure changes of GRXS17 and GRXS17-C33/179/309/416S with circular dichroism spectroscopy (Supplemental Figure 2). The general structures (α-helices, β-strands and turn) remains similar between both proteins, indicating that the C33/179/309/416S mutations does not induce major secondary structure changes of GRXS17. The effect of Cys oxidation on the conformation of GRXS17 was also determined. Treatment with 400 µM H_2_O_2_ resulted in a minor change in secondary structure composition of wild-type GRXS17, but not for the GRXS17-C33/179/309/416S variant (Supplemental Figure 2A). This minor change in secondary structure composition of GRXS17 upon oxidation can correspond to either a local or a global conformational change, which is likely to involve the partial unfolding of α-helices and the formation of β-strands. As this conformational change was not observed for the quadruple Cys variant GRXS17-C33/179/309/416S, we can conclude that this structural change depends on the oxidation of the catalytic cysteines and the formation of disulfide bonds in wild-type GRXS17.

To summarize, we showed that recombinant GRXS17 is able to oligomerize by forming intermolecular disulfides in which the active site C33 of the TRX-domain plays a major role. In addition to being involved in Fe-S cluster coordination, C179, C309 and C416 seems to be important for maintaining non-covalent interactions between protomers. Finally, the active site residues C33, C179, C309 and C416 are not involved in the dimerization process.

### A holo-apo redox switch of GRXS17 activates holdase activity upon oxidation

Molecular chaperone activities are often associated with oligomerization and are related to thermotolerance (Lee et al., 2009). Therefore, the potential role of GRXS17 as a molecular chaperone was analyzed using different chaperone substrates. In order to monitor holdase activity, we used mitochondrial citrate synthase (CS) as a client protein (Figure 3). We analyzed the effect of GRXS17 on the high-temperature-induced aggregation of CS, a classical assay to monitor the activity of molecular chaperones *in vitro* (Voth et al., 2014). Incubation of CS with reduced or oxidized apo-GRXS17 prevented thermal aggregation at 43°C (Figure 3A). Increasing the apo-GRXS17/CS ratio progressively decreased CS aggregation which was fully prevented at a molar ratio of 0.5:1 apo-GRXS17:CS (Figure 3B). This effect was stronger than described for holdases from various organisms (e.g., HSP33, Get3, CnoX, TDX) (Jakob et al., 1999; Lee et al., 2009; Voth et al., 2014; Goemans et al., 2018). In contrast, in the absence of GRXS17 or in the presence of an excess of ovalbumin, CS significantly aggregates after 15 min (Figure 3A). Interestingly, comparable holo-GRXS17/CS ratios did not prevent CS aggregation, indicating that the holdase activity of GRXS17 is specifically associated with the apo-GRXS17 form (Figure 3A). Subsequently, we verified that the Fe-S cluster in the holo-GRXS17 is stable at 43°C (Supplemental Figure 1).

**Figure 3:**
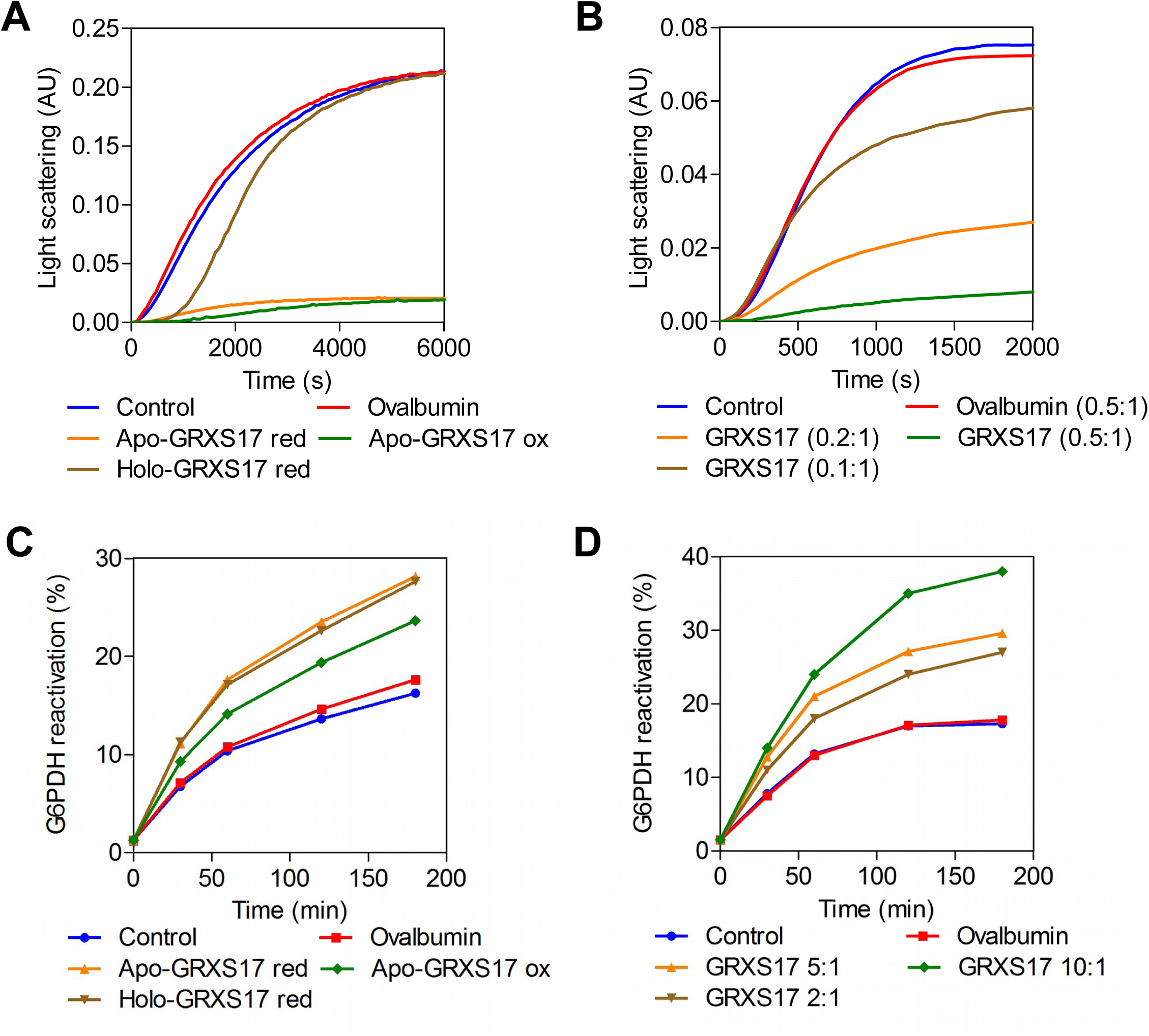
GRXS17 exhibits holdase and foldase chaperone activities. **(A-B)** GRXS17 holdase activity was measured towards citrate synthase (CS) and malate dehydrogenase (MDH) by light scattering. **(A)** Holo-GRXS17 or apo-GRXS17 in their reduced or oxidized forms were incubated with CS at a 0.5:1 ratio at 43°C. Apo-GRXS17 but not holo-GRXS17 protects CS from temperature-induced precipitation. **(B)** Different ratios of apo-GRXS17/MDH were incubated at 55°C. Similar results were found by incubating apo-GRXS17/CS at 43°C. **(C-D)** GRXS17 foldase activity was measured by reactivation of the glucose 6-phosphate dehydrogenase (G6PDH). **(C)** Holo-GRXS17 or apo-GRXS17 in their reduced or oxidized forms were incubated with G6PDH at a 5:1 ratio at 30°C. Apo-GRXS17 and holo-GRXS17 support reactivation of G6PDH. **(D)** The ratio of apo-GRXS17/G6PDH was varied. As controls, substrates were incubated alone (Control) or in presence of ovalbumin. Data are representative of three different experiments.

We also monitored the foldase activity of GRXS17 by using glucose-6-phosphate dehydrogenase (G6PDH) as a substrate. To distinguish between foldase and disulfide reductase activities, we used urea-denatured Cys-free G6PDH. Increasing the apo-GRXS17:G6PDH ratio progressively restored the G6PDH activity (Figure 3C-D). This reactivation was significantly faster in the presence of GRXS17 with nearly 36% recovery after 3 h at a 10:1 ratio of GRXS17:G6PDH. Reduced or oxidized holo-GRXS17 similarly restored G6PDH activity, indicating that the foldase activity of GRXS17 is independent of the presence or absence of the Fe-S cluster (Figure 3C-D). We also could show that these activities are ATP-independent (Supplemental Figure 3). Collectively, our data show that GRXS17 can adopt both a holdase and a foldase activity *in vitro,* and that the holdase activity is specific for the apo-form.

### Oligomerisation determines the holdase activity of GRXS17

The capacity of apo-GRXS17 to oligomerize depends on both disulfide bonds involving active site Cys residues and non-covalent electrostatic interactions (Figure 2). In order to uncover forms that are associated with chaperone activity, we tested the chaperone activity of the Cys/Ser-variants of GRXS17 using cytosolic malate dehydrogenase (cytMDH1) as substrate (Figure 4A). We found that GRXS17(C33S) has holdase activity for those forms with a high oligomerization state (Figure 2C and Figure 4A). In the oligomerization-abolished GRXS17-C33/179/309/416S variant (Figure 2D) no holdase activity was observed (Figure 4A). This suggested that the holdase activity depends on the active-site cysteines. To further demonstrate that oligomerization is required for holdase activity, we incubated reduced and oxidized apo-GRXS17 with a low concentration of the chaotropic agent guanidinium hydrochloride (GdnHCl), which abolishes weak-bonds like hydrophobic and electrostatic interactions (Figure 4B). We found that the holdase activity of reduced GRXS17 is inhibited with GdnHCl, while the holdase activity of oxidized GRXS17 is preserved. Finally, we tested the role of the cysteines in GRXS17 foldase activity (Figure 4C). We found that GRXS17 Cys mutations does not negatively affect GRXS17 foldase activity, suggesting that disulfide-linked oligomerization is not required for its foldase activity.

**Figure 4:**
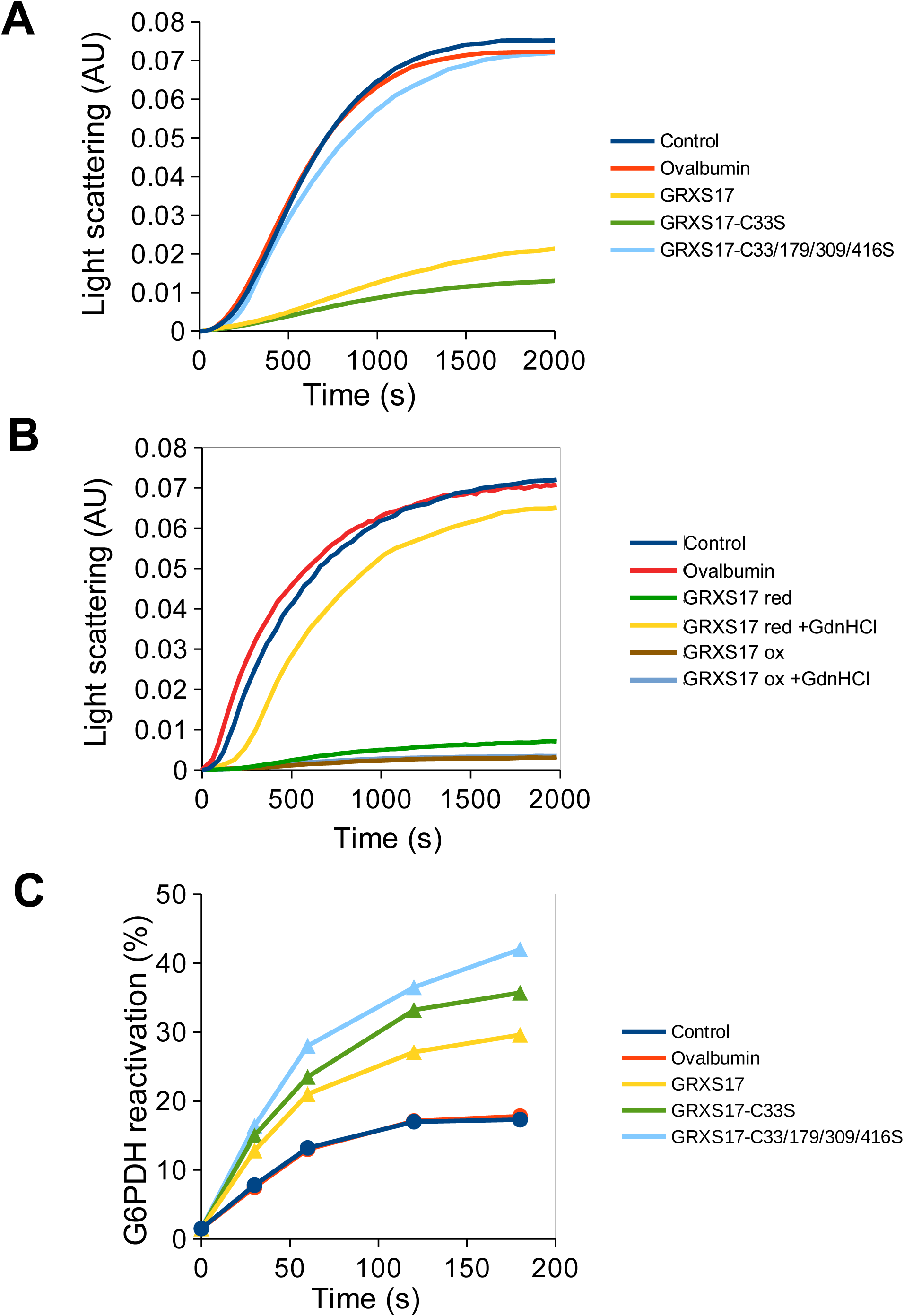
Role of GRXS17 active-site cysteines and oligomerization in holdase and foldase activities. **(A)** GRXS17 and Cys-mutated GRXS17 proteins holdase activity was measured towards cytMDH1 heat-induced precipitation by light scattering at 0.5:1 ratios at 55°C. CytMDH1 was incubated alone (Control), with ovalbumin or with GRXS17 proteins. **(B)** GRXS17 proteins reduced by 5 mM DTT (green) or oxidized with 1 mM H_2_O_2_ (brown) in absence or presence (respectively yellow and light blue) of 20 mM guanidinium hydrochloride (GdnHCl) as chaotropic reagent. **(C)** GRXS17 foldase activity was determined by measuring the glucose-6 phosphate dehydrogenase (G6PDH) activity after denaturation with urea at a 5:1 ratio. CytMDH1 was incubated alone (Control), with ovalbumin or with GRXS17 proteins. Data are representative of three independent experiments.

### GRXS17 is involved in tolerance to moderate heat stress

To reveal the physiological significance of the redox-dependent chaperone activity of GRXS17, we explored the possible role of GRXS17 in plant thermotolerance. Therefore, we subjected *grxs17* mutant plants to different heat-stress regimes (Yeh et al., 2012). The impact of heat stress on shoot meristem viability was first quantified by monitoring the appearance of new leaves. The viability of the *grxs17* mutant was similar to wild-type plants for both Short-term Acquired Thermotolerance (SAT) and Long-term Acquired Thermotolerance (LAT) regimes (Figure 5A-B). In contrast, a mutant lacking the heat-shock protein 101 (HSP101) chaperone was found to be hypersensitive to all heat-stress regimes (Yeh et al., 2012), indicating that GRXS17 does not overlap with HSP101 functions in SAT and LAT. Nevertheless, compared to wild-type plants, *grxs17* mutants were significantly more sensitive to a moderate heat stress regime (TMHT: Tolerance to Moderated High Temperature) (Figure 5C), indicating that GRXS17 is specifically involved in this heat stress regime. The thermosensitive phenotype of *grxs17* was fully complemented by overexpressing the full-length GRXS17:GFP fusion, indicating that the TMHT phenotype of *grxs17* is due to the loss of the GRXS17 function and that the attached GFP does not perturb the function of the protein (Figure 5C and Supplemental Figures 4 and 5). Importantly, the overexpression of the GRXS17-C33S:GFP fusion was able to restore the wild-type phenotype, while the expression of the GRXS17-C33/179/309/416S:GFP variant failed to complement the mutant phenotype (Figure 5C, Supplemental Figures 4 and 5). This last observation demonstrates the role of active site Cys in moderate heat stress tolerance.

**Figure 5:**
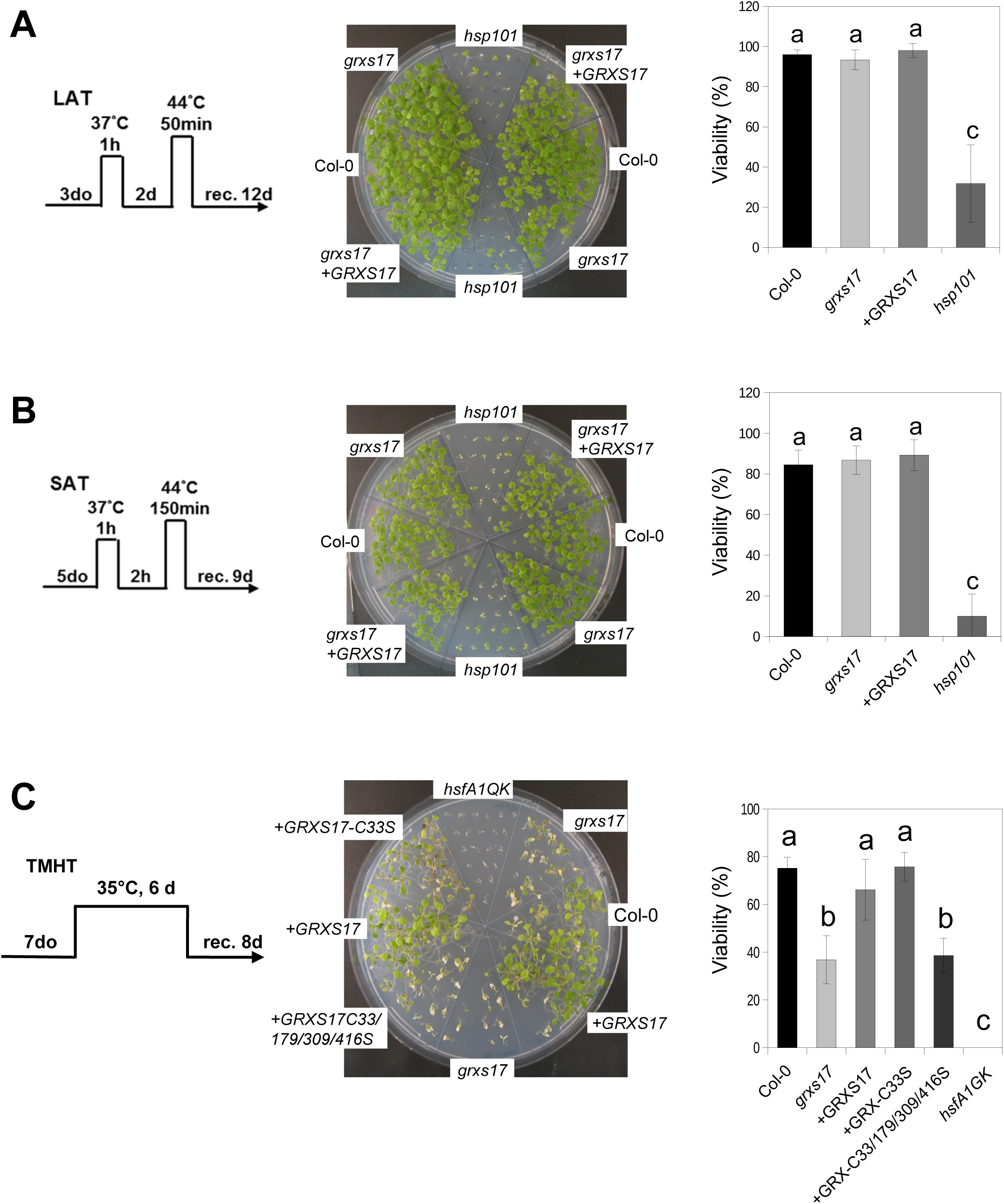
GRXS17 is specifically involved in Thermotolerance to Moderated High Temperature. Plants were subjected to different temperature stress regimes. (**A**) LAT, Long-term Acquired Thermotolerance. **(B)** SAT, Short-term Acquired Thermotolerance). **(C)** Thermotolerance to Moderated High Temperature (TMHT) experiment. do: days old. rec: recovery. Wild-type plants (Col-0), Heat-Shock Factor A1 quadruple knock-out mutant (*hsfA1QK*), Heat-Shock Protein 101 knock-out mutant (*hsp101*), *grxs17* knock-out mutant (*grxs17*), *grxs17* mutant complemented with a *Pr35S:GRXS17:GFP* construct expressing the GRXS17:GFP fusion protein (+GRXS17), the GRXS17-C33S:GFP protein (+GRXS17-C33S) or the GRXS17-C33/179/309/416S:GFP (+GRXS17-C33/179/309/416S) protein. Viability of the plants was assessed by recovery of the shoot growth. Data are means of 4-10 biological repetitions +/- SE, n = 20-25. The letters a, b and c indicate a significance of p>0.1, p<0.001, and p<0.00001 respectively, by Student’s t-test.

In order to study the impact of high temperature on root, wee performed a similar experiment by following the root development after the TMHT regime (Figure 6). Plants were grown at 20°C for 6 days before the temperature was shifted to 30°C. For root growth, 30°C was used instead of 35°C because this latter temperature affected to much root growth of the wild-type plants (not shown). We observed that growth of the primary root of wild-type seedlings tends to be accelerated after the temperature shift, while root growth of the *grxs17* mutant almost completely stopped, confirming the sensitivity of the mutant to moderate heat stress. We also observed a dramatic decrease of emerged lateral root number in the *grxs17* (Figure 6B). When the *grxs17* mutant was complemented with the GRXS17-GFP fusion protein, the mutant behaves like wild-type. On the other hand, GRXS17-C33/179/309/416S:GFP fails to complement the *grxs17* mutant (Figure 6A-B). The structure of the root apical meristem (RAM) was observed at 11 days of growth (after 5 days of heat stress). Consistently, the *grxs17* mutant presented a significantly shorter RAM compared to wild-type and the GRXS17-GFP complemented lines, showing a high impact of heat stress on the meristematic activity of the mutant (Figure 6C-E). Propidium iodide (PI) staining was used to monitor cell death in meristematic zones (Truernit and Haseloff, 2008). While we did not observe any cell death in the RAM, massive cell death was observed in the lateral root primordia in *grxs17* (Figure 6G-I). This suggested that GRXS17 prevents heat stress injuries at an early stage of meristem establishment, which we confirmed in a germination experiment under heat stress using the *grxs17* mutant. While the mutant seeds are able to germinate properly, they stop growing at early plantlet stage and harbor a very short root. Importantly, the structure of the RAM looks disorganized and shows extensive cell death (Supplemental Figure 6). The GRXS17-C33/179/309/416S:GFP complemented lines failed to complement the shorter RAM of the *grxs17* mutant and did not prevent cell death in the lateral root initials, indicating the key role of Cys in GRXS17 functions (Figure 6F and J). In addition, we also observed perturbations in the root hair morphology in these latter plants, which were not observed in any other conditions, suggesting a deleterious effect of GRXS17-C33/179/309/416S:GFP overexpression in root hairs (Figure 6J, inset).

**Figure 6:**
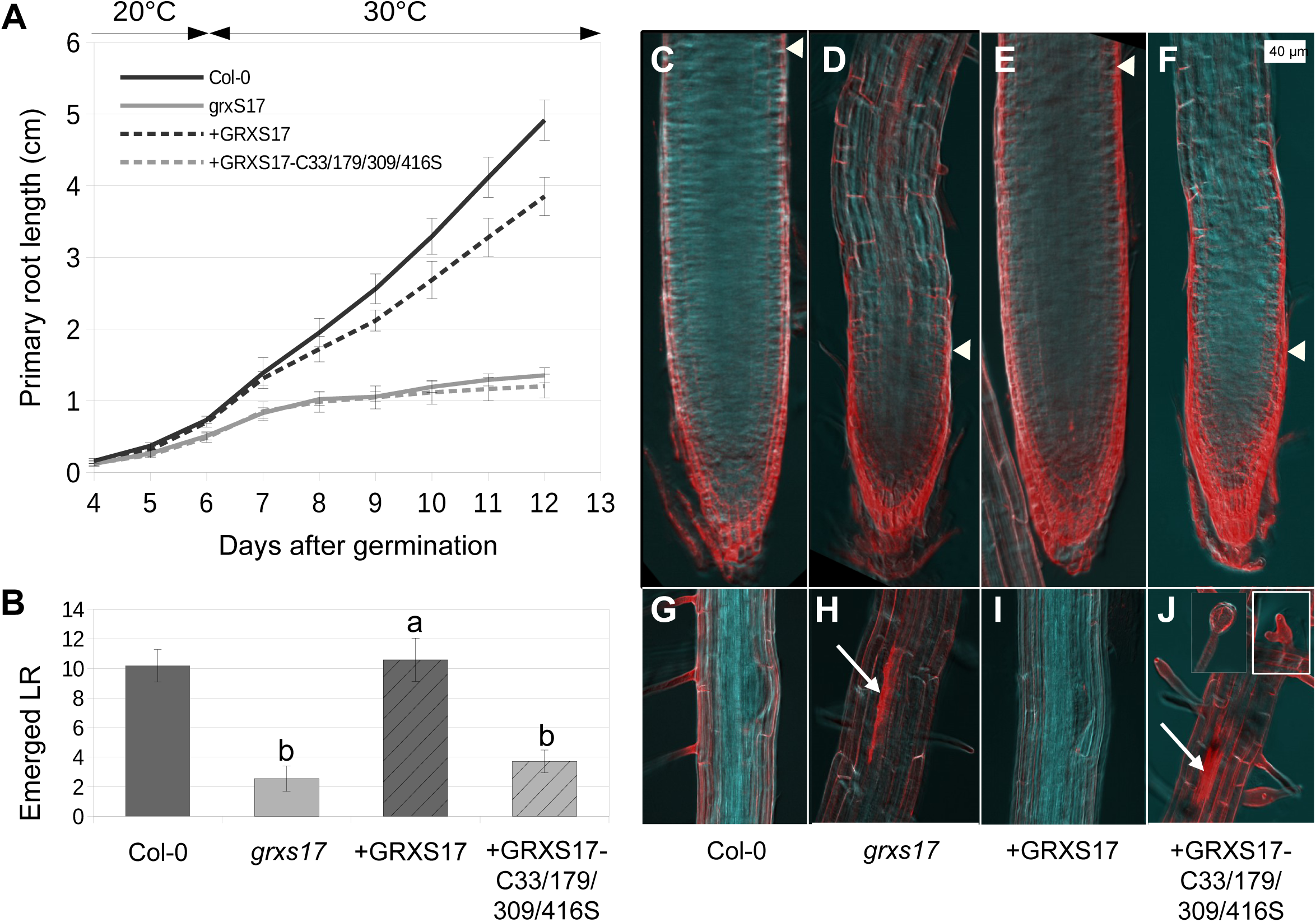
GRXS17 is involved in root development under heat stress. Plants grown 6 days at 20°C were shifted to 30°C for 6 more days. **(A)** Primary root length. **(B)** Number of emerged lateral roots in 12 day-old plants. **(C-F)** Primary root meristem and **(G-J)** lateral root primordia in 12 day-old plants stained by propidium iodide. Wild-type plants (Col-0), *grxs17* knock-out mutant (*grxs17*), *grxs17* mutant complemented with a *Pr35S:GRXS17:GFP* construct (+GRXS17) and the *Pr35S:GRXS17-C33/179/309/416S:GFP* construct (+GRXS17-C33/179/309/416S). White triangles indicate the limit between the meristematic and the elongation zone. White arrows indicate dead cell in root primordia. Insets represent ectopic morphology of root hair specifically observed in the complemented line +GRXS17-C33/179/309/416S. Data are means of 3 biological repetitions +/- SE, n = 20. The letters a and b indicate a significance of p>0.1 and p<0.0001 respectively, by Student’s t-test.

Altogether, these results highlight the role of GRXS17 in the tolerance of plants to moderate heat stress, acting to protect both shoot and root meristems. Importantly, the active site cysteines of GRXS17 seems to play an essential role in root growth and thermotolerance.

### GRXS17 oligomerizes *in planta* and associates with a different set of proteins upon shifting to a higher temperature

In addition to *in vitro* assays, we also explored GRXS17 oligomerization *in vivo*. Total soluble protein extracts prepared from Arabidopsis seedlings grown at standard temperature (20°C) and exposed to heat stress (35°C for 2 h) were fractionated by SEC (Figure 7). Western-blot analyses of eluted fractions revealed a peak containing GRXS17 eluting at ∼100–150 kDa (lanes 23–25), corresponding to a molecular weight of a GRXS17 dimeric form (∼110 kDa) (Knuesting et al., 2015). Moreover, at 20°C, a GRXS17 signal was also detected in fractions (lanes 18–21) corresponding to an HMW (∼380 kDa) form (Figure 7). Interestingly, plants subjected to heat stress accumulated higher levels of HMW forms eluting in the range from SEC column (lanes 16-18), corresponding to 520 kDa (Figure 7). These data suggest that similarly to *in vitro* analyses (Figure 1 and 2), GRXS17 forms, HMW-complexes *in vivo*, although the size of these complexes are not fully similar. Also, we cannot rule out that GRXS17 oligomeres associate with other proteins.

**Figure 7:**
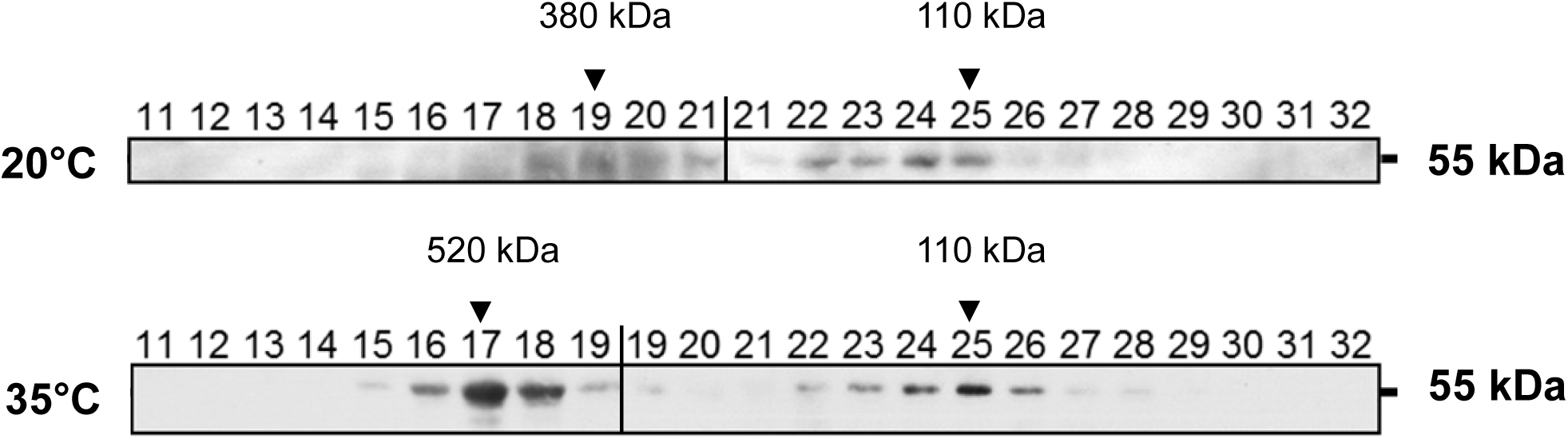
GRXS17 forms high molecular weight complexes in plant extracts. Size-exclusion chromatography analysis (Sephacryl S300 HR) of crude protein extracts from Arabidopsis seedlings cultivated 14 days at 20°C and further treated for 2 h at 35°C. The GRXS17 protein was detected after western bloting and immunodetection using an anti-GRXS17 serum. Numbered lines correspond to the protein fractions. Protein size of the respective fractions was obtained after calibration using marker proteins.

Finally, we sought to identify GRXS17 partners by using immunoprecipitation coupled to mass-spectrometry determination (IP-MS). To this end, *grxs17* mutant plants were transformed with a *Pr35S::GRXS17:FLAG-HA* construct. The expression of the construct was verified by immunoblotting and its functionality was assessed by complementation of the phenotype under TMHT (Supplemental Figure 7). Total protein extracts from GRXS17:FLAG-HA expressing plants grown at 20°C or shifted for 2 h to 35°C were collected for IP-MS analyses. Non-specific GRXS17 contaminants were identified within the same experiments using *grxs17* mutant plants expressing a GRXS17 protein devoid of the FLAG-tag. We employed a label-free quantitative proteomic approach to identify proteins significantly enriched after IP by comparison with the control samples. Five biological replicates were analyzed by mass spectrometry after FLAG immunoprecipitation from total protein extracts of plants expressing GRXS17:FLAG-HA or non-tagged GRXS17 (control) grown at 20°C or shifted for 2 h to 35°C. GRXS17 was the only protein significantly enriched in the tagged samples after IP (P<0.05, Bonferroni corrected Student T-test). In addition, we identified 23 proteins from plants grown at 20°C and 12 proteins from plants shifted to 35°C that were present and quantified in at least 3 of the 5 replicates from the tagged strain and not found in any of the 5 replicates from control non-tagged plants (Table 1). Using this all-or-nothing approach, several likely interactants such as protein disulfide isomerase (PDIL) and the antioxidant enzyme monodehydroascorbate reductase (MDAR) were identified at 20°C. A different set of interaction partners was identified after a 2 h temperature shift from 20°C to 35°C, and only 4 proteins were identified in both conditions (Table 1). Among them, the nucleo-cytoplasmic BolA2 was always present (Supplemental Table 1). This interaction is consistent with previous studies that reported the interaction of GRXS17 with BolA2 (Couturier et al., 2014). Finally, to identify

**Table 1:**
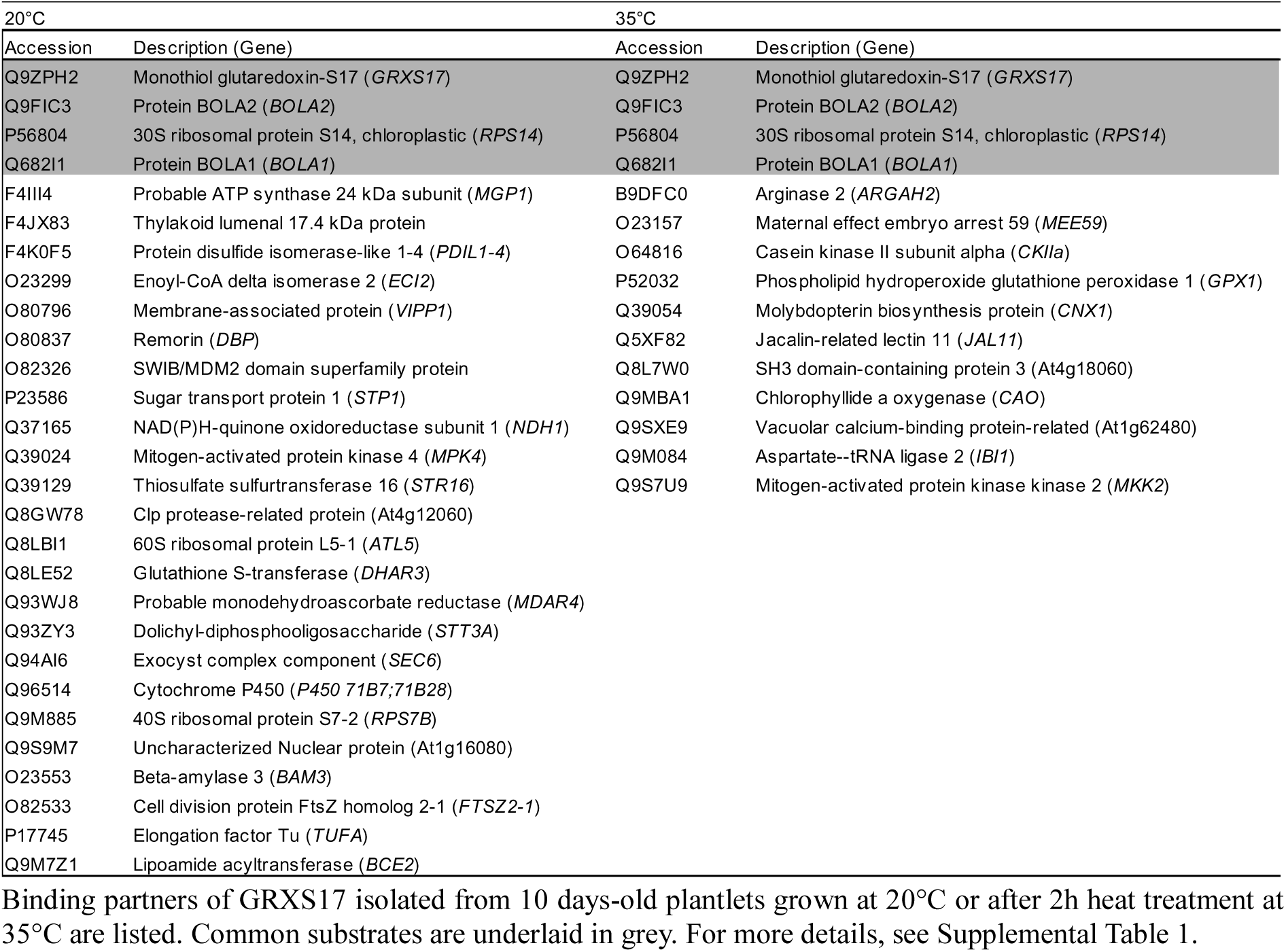
Proteins interacting with GRXS17 at 20°C or 35°C

GRXS17 partners in HMW complexes, total protein extracts from plants grown at 20°C or shifted for 2 h to 35°C were first separated by SEC, and HMW fractions containing GRXS17:FLAG-HA were collected for IP-MS analysis. However, a simple subtractive analysis between the samples failed to identify a similar set of proteins as in the previous approach, suggesting that those proteins are not involved in HMW complexes (Supplemental Table 1). However, BolA2 was identified after IP-MS in HMW fractions, indicating that BolA2 is a major GRXS17 oligomer interactant associated with the molecular chaperone activity of GRXS17.

## DISCUSSION

### Apo-GRXS17 switches to holdase activity upon stress

During periods of heat stress, living organisms need to rapidly adapt to cope with the damaging effects of increasing temperatures. The sessile nature of plants makes them particularly vulnerable to heat stress. As in other organisms also in plants, a set of heat-shock proteins acts as molecular chaperones to prevent damage caused by protein inactivation and aggregation. Most functions of heat-shock proteins are heat-inducible, but their activation mechanism is not well understood (Hartl et al., 2011). Here, we demonstrated that the Fe-S cluster-harboring GRXS17 becomes a holdase after exposure to heat in oxidizing conditions. This switch happens within minutes by a redox and temperature destabilization of the Fe-S cluster, and subsequent disulfide bond formation (Figure 8). The redox-switch of Fe-S cluster-containing GRX has previously been shown to occur under oxidizing conditions in other GRX. In these cases, the active site Cys are involved in the Fe-S cluster which inhibits the thiol-reducing activity, the loss of the Fe-S cluster restoring the activity (Lillig et al., 2005; Berndt et al., 2007; Rouhier et al., 2007; Bandyopadhyay et al., 2008; Riondet et al., 2012). In contrast, the apo-GRXS17 of this study is inactive as a thiol reductase, as also reported for another class-II GRX (Bandyopadhyay et al., 2008; Knuesting et al., 2015; Moseler et al., 2015). This lack of thiol reductase activity was proposed to be linked to the specificity of the active site and the low reactivity for GSH (Moseler et al., 2015; Begas et al., 2017).

**Figure 8:**
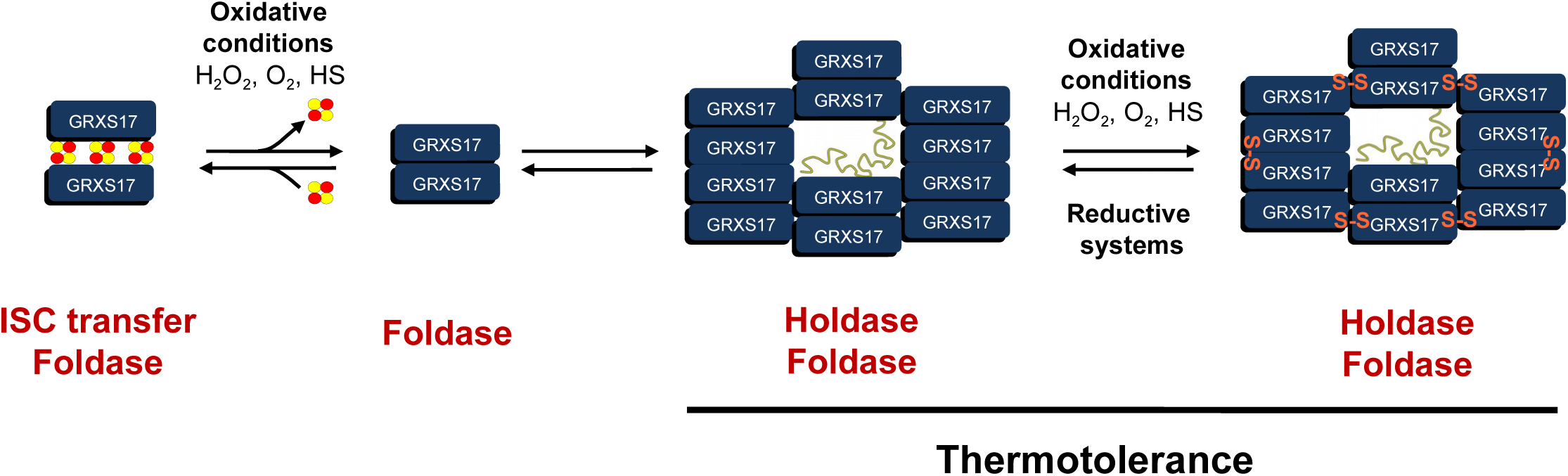
Model of the GRXS17-redox switch. Under reducing conditions, the holo-GRXS17 forms a dimer which coordinates an iron-sulfur cluster (ISC). In this form, GRXS17 can transfer ISC to cytosolic and nuclear target proteins (e.g., BolA). Under oxidative conditions (H_2_O_2_, O_2_, Heat Stress (HS)), ISC are released and GRXS17 switches to an apo-dimeric form with a foldase activity. Dimers can spontaneously associate to oligomeric forms and further oxidation induces oligomeric structure through formation of intermolecular disulfide bonds at active-site Cys residues. In this form, GRXS17 acquires holdase activities and provides thermotolerance. The oligomeric form can be reduced to recover the apo-dimeric GRXS17 and the Fe-S cluster is reassembled.

Interestingly, the active-site cysteines of GRXS17 are crucial for the holdase redox switch as they coordinate both the Fe-S cluster, and respond to an oxidizing environment, such as aerobic conditions, H_2_O_2_ exposure, or heat stress. As such, GRXS17 may act as a sensor for cellular redox changes. In fact, high temperatures by itself do not destabilize the Fe-S cluster, but the simultaneous presence of H_2_O_2_ oxidatively changes the conformation of the protein and induces oligomerization through non-covalent interactions and disulfides (Figure 8). These observations indicate that the Fe-S cluster stabilizes the dimeric structure of GRXS17, and only after dissociation the protein can undergo oligomerization. A similar function has previously been shown in Archaea, where the loss of a [4Fe-4S] cluster causes oligomerization of the D subunit of the RNA polymerase (Hirata et al., 2008). In GRXS17, we clearly showed that the active-site cysteines (C33, C179, C309 and C416) are required for both non-covalently and covalently mediated oligomerization (Figure 2). Although GRXS17 was shown to reduce *in vitro* a monomeric glutathionylated form and a dimeric disulfide-bridged form of BolA2 (Couturier et al., 2014), the oligomerization of the apo-GRXS17 protein under oxidative conditions may inhibit this activity (Ströher et al., 2016).

Proteins that combine chaperone function with a redox activity previously have been described. For example, plant thiol reductase proteins TRXh3, TDX and NTRC exhibit chaperone activities triggered by oxidative stress and heat shock exposure, leading to a shift of the protein structure from a low-MW form to high-MW complexes (Lee et al., 2009; Chae et al., 2013; Park et al., 2009). Also, overoxidation of nucleophilic cysteine of 2-Cys PRX triggers oligomerization and induces chaperone activity (Liebthal et al., 2018). GRXS17 is different as it has no oxidoreductase activity, but it has a much more efficient holdase activity than the thiol oxidoreductases (Figure 3 and Lee et al., 2009; Chae et al., 2013; Park et al., 2009). Remarkably, the key to induce holdase chaperone activity is the release of the redox-dependent Fe-S cluster. To our knowledge, this is the first example of an FeS protein that turns into a chaperone upon switching between holo- and apo-form. In *E. coli*, a similar example is hsp33 which coordinates zinc via conserved cysteines (Jakob et al., 1999; Jakob et al., 2000; Winter et al., 2008). Upon oxidation with H_2_O_2_, disulfide bonds are formed between the coordinating cysteines and zinc is released, activating HSP33 holdase activity. Here, zinc coordination, oxidation and activation state of hsp33 are directly linked to the redox state of the environment (Jakob et al., 1999; Jakob et al., 2000; Winter et al., 2008). As another example in yeast, the cytosolic ATPase Get3 turns into an effective ATP-independent chaperone when oxidized (Voth et al., 2014). Furthermore, the holdase activity of *E. coli* chaperedoxin CnoX is activated by chlorination upon bleach treatment to protect substrates from irreversible oxidation (Goemans et al., 2018). Whether GRXS17 acts by protecting its interactors from overoxidation remains to be determined.

### GRXS17 exhibits a constitutive foldase activity

GRXS17 is likely acting as a housekeeping chaperone involved in folding cytosolic (or nuclear) nascent peptides or in refolding misfolded peptides. We found that the foldase activity of GRXS17 is independent of ATP (Supplemental Figure 3). This clearly differs from most prokaryotic (e.g. GroEL-GroES) and eukaryotic foldases (e.g. hsp70, hsp90) which need ATP for their foldase activity (Mayer, 2010; Hartl et al., 2011). Nevertheless, a number of chaperones have been identified that promote folding in the absence of high-energy cofactors (Stull et al., 2016; Horowitz et al., 2018). As an example, the small ATP-independent chaperone Spy from *E. coli* uses long-range electrostatic interactions and short-range hydrophobic interactions to bind to its unfolded client proteins and drive their folding (Koldewey et al., 2016).

Protein disulfide isomerase (PDI), or cyclophilin, which catalyze disulfide formation and isomerization of disulfides, exhibit folding activities through their active site Cys (Park et al., 2013; Ali Khan and Mutus, 2014). GRXS17 functions differently, as the foldase activity of GRXS17 is independent of its active-site cysteines (Figure 4). Nevertheless, two additional cysteines are present in the first and third GRXS17 domains, and their involvement in GRXS17-foldase activity cannot be ruled out. Future experiments will have to be performed to decipher the mechanistic of the foldase activity of GRXS17.

### GRXS17 associates with different proteins under high temperature

In eukaryotic organisms grown under non-stress conditions, the cytosolic compartment is a highly reducing environment (Schwarzländer et al., 2008). However, in heat-stressed plants, intracellular oxidation triggered by ROS accumulation was largely documented (Choudhury et al., 2017; Dickinson et al., 2018). This is consistent with measurements of the glutathione redox potential (*E_GSH_*) monitored in plant expressing the GRX1-roGFP2 construct in the cytosol and nucleus and subjected to high temperature (Supplemental Figure 8). Therefore, in regard to changes in redox status depending on the temperature, it is likely that the major GRXS17 form is the Fe-S cluster-containing dimeric form under standard growth temperatures, but a significant proportion of GRXS17 switches to HMW complexes in plants subjected to heat stress (Figure 7). Although the difference in the size of the HMW complexes observed in recombinant GRXS17 and in plant extract will have to be further explored, the switch to HMW complexes appears to be necessary for plant survival under heat stress. As the *grxs17* mutant is highly sensitive to moderate high temperature regimes, the mutant can only be complemented by the GRXS17-Cys variants that are still able to oligomerize (GRXS17 and GRXS17-C33S) and not by GRXS17-C33/179/309/416S (Figures 5 and 6) (Cheng et al., 2011; Knuesting et al., 2015). Further, this demonstrates the crucial role of the active site Cys in the thermosensitive function of GRXS17. The role of Fe-S cluster in thermotolerance will have to be further studied by generating mutant *grxs17* plants complemented by the GRXS17-C179/309/416S in which only cysteines involved in Fe-S cluster coordination (and not in chaperone activity and oligomerization) are mutated. We unfortunately failed to generate such plants yet.

GRXS17 was previously proposed to improve thermo-, drought- and oxidative-tolerance by enhancing ROS scavenging capacities and expression of Heat-Shock Proteins. This might occur by modulation of gene expression, enzyme activity or protection of antioxidant enzymes (Wu et al., 2012, 2017; Hu et al., 2017). Our data suggest an additional function of GRXS17 through its chaperone activity. The GRXS17 partner protein identification resulted in a low overlap between proteins immunoprecipitated at 20°C and those identified at a higher temperature, suggesting that the set of interacting proteins is completely changed under heat stress. Although it is tempting to suggest that the proteins we identified in immunoprecipitation experiments are protected by GRXS17, further experiments will need to validate these partners and the biological significance of these interactions. Importantly, our observations showing that GRXS17 maintains plant viability under moderate heat stress conditions and prevents cell death in founder cells of root primordia under high temperature might indicate a role in protecting key actors of meristem function. Possible candidates that we found in our IP are MPK4 and MKK2 which play roles in temperature signalling pathways (Zhao et al., 2017).

Finally, we propose that the unique properties of GRXS17 among the redox-regulated chaperones (e.g., 2-Cys PRX, TRXh3 or TDX) comes from its inducible holdase function (i.e. Fe-S cluster release under stress), which is not seen for other heat stress-related chaperones (Stull et al., 2016). Homologs of GRXS17 are present in all plants and in most other organisms (Couturier et al., 2009), and their functions in iron metabolism are well documented (Encinar del Dedo et al., 2015; Berndt and Lillig, 2017). It is likely that the switch to holdase chaperone activity, which we described here, could be conserved among different kingdoms of life. Thus, unlike other Fe-S cluster-containing proteins in which cluster release generally impairs protein function (Lill, 2009; Klinge et al., 2007; Berndt and Lillig, 2017), GRXS17 uses this oxidation-induced switch to activate holdase functionality, and in this way, protects plants from heat-induced damage.

## METHODS

### Plant materials and growth conditions

Wild type (WT) *Arabidopsis thaliana* ecotype Columbia-0 (Col-0), *grxs17* (SALK_021301) mutants were already characterized (Knuesting et al., 2015). Arabidopsis seeds were surface-sterilized for 15 min with ethanol 70%, rinsed with ethanol 95% and completely dried before used. Sterilized seeds were plated onto ½ MS agar medium (2.2 g/L Murashige and Skoog, 0.5 g/L MES, 1% plant agar (w/v), adjusted to pH 5.8) and stratified 48 h at 4°C in the dark. Seedling were grown under 120 µmol.m^-2^.s^-1^ photosynthetic flux at 22°C and 16 h light/8 h dark cycles. Those growth conditions were systematically used, unless otherwise indicated.

### Gene cloning, mutagenesis, plasmid construction

For the construction of transgenic complemented line of the *grxs17* mutant and production of recombinant proteins, cDNA of GRXS17 (At4g04950) were inserted into the pGEM-T Easy plasmid (Promega, Madison, WI, USA). Different point mutations of cysteines to serines in GRXS17 were generated using QuikChange II Directed Mutagenesis Kit (Agilent) using primers detailed in Supplemental Table 2: GRXS17-C33S (GRXS17-m1), GRXS17-C179S/C309S/C416S (GRXS17-m234) and GRXS17-C33S/C179S/C309S/C416S (GRXS17-m1234).

To generate complemented lines, *GRXS17, GRXS17-C33S, and GRXS17-C33S/C179S/C309S/C416S* genes were cloned in the pCAMBIA1302 (*Pr35S:gene:GFP*) vector. To perform immunoprecipitation, *GRXS17* was cloned in a modified pCAMBIA 1300 vector harboring a Flag-Flag-HA-HA sequence fused to the C-terminal end of the protein (*Pr35S:gene:Flag/Flag/HA/HA*).

For the production of the recombinant proteins, GRXS17, GRXS17-C33S, GRXS17-C179/309/416S, GRXS17-C33/179/309/416S and the cytosolic malate dehydrogenase1 (cytMDH1) from *Arabidopsis thaliana* (At1g04410) were cloned in the pET16b vector (Novagen; Merck Biosciences) using primers detailed in Table 1.

### Transformation of plants

Plasmids were transferred into Arabidopsis via *Agrobacterium tumefaciens* strain GV3101 using the described methods (Clough and Bent, 1998). Transformants were selected on ½ MS medium plates containing 50 μg ml^-1^ hygromycin and checked by PCR using appropriate primers (Supplemental Table 2).

### Production and purification of recombinant proteins

All proteins were produced by induction in *Escherichia coli* BL21(DE3). Cultures of 2 L of a ampicillin-resistant (100 mg.mL^−1^) colony were grown at 37 °C and induced by 100 µM isopropyl-β-D-galactopyranoside (IPTG) in the exponential phase (DO=0.5). Bacteria were harvested by centrifugation 3 h after induction and stored at -80°C before protein extraction. Pellets were resuspended in buffer (50mM Tris-HCl pH 7.0, 100 mM NaCl, and 1 tablet anti-protease cocktail (Roche) for 25 mL), extracted using a cell disrupter (Constant Systems Limited, Northants, UK) and purified with a commercial kit according to manufacturer’s instructions (His-bind buffer kit, Millipore).

### *In vitro* characterization of holo-GRXS17

Chemical reconstitution of holo-GRXS17 was performed under strictly anaerobic conditions in an anaerobic vinyl tent (COY Laboratory Products, Grass Lake, U.S.A.) according to Freibert et al. (2018). In brief, 100 μM GRXS17 were reduced in reconstitution buffer (50 mM Tris-HCl, pH 8, 150 mM NaCl, 10% glycerol (v/v)) in the presence of 2 mM GSH and 2 mM DTT for 3 h at room temperature. For reconstitution, the GRXS17 concentration was diluted to 20 μM and ferric ammonium citrate was added to a final concentration of 400 μM. After 5 min of incubation, lithium sulfide (400 μM)) were added followed by a 3-h incubation. Excess iron ammonium citrate and lithium sulfide was removed by desalting using a NAP™ 25 column (GE Healthcare). Subsequently, the ISC were characterized using a CD-spectrophotometer (J-815, Jasco). The reconstituted GRXS17 was transferred to a CD cuvette, which was sealed airtight. Oxidative treatment of the holo-GRXS17 was performed with air oxygen (21%) for the indicated time. The analytical SEC was carried out under strict anaerobic conditions. The gel filtration was performed with the ÄKTA Prime Plus System (GE Healthcare) coupled to a Superdex200 10/300 GL column (GE Healthcare). In each case, 50 μg of GRXS17 were applied into the equilibrated column (50 mM Tris-HCl pH 8, 150 mM NaCl, 10% glycerol (v/v)) and separated at a flow rate of 0.5 mL/min. The column was calibrated with the Molecular Weight standard 29-669 kDa (Sigma-Aldrich).

### Holdase and foldase activities

The holdase chaperone was measured by using the cytosolic MDH1 (cytMDH1) from *Arabidopsis thaliana* (Huang et al., 2018) or the citrate synthase (CS) from porcine heart (Sigma-Aldrich) at 55°C or 43°C, respectively according to Lee et al. (2009). The foldase chaperone activity was examined by using glucose-6-phosphate dehydrogenase (G6PDH) (Sigma-Aldrich) according to Lee et al. (2009). Light scattering (holdase) or absorption (foldase) were monitored with a Shimadzu UV-1800 spectrophotometer at 340 nm (Shimadzu) equipped with a thermostated cell holder.

### Confocal Laser-Scanning Microscopy and cell death observation

Confocal microscopic observations were carried out using the Axio observer Z1 microscope with the LSM 700 scanning module and the ZEN 2010 software (Zeiss). GFP and propidium iodide (PI) were excited using the 488nm argon-ion laser and collected at 500-550 nm (GFP) or 600-656 nm (PI). Excitation of roGFP2 was performed at 488 and 405 nm and a bandpass (BP 490-555 nm) emission filter was used to collect roGFP2 signal. For background subtraction, signal was recovered using a BP 420-480 nm emission filter during excitation at 405 nm. Pictures analyses and quantifications were performed as previously described (Schwarzländer et al., 2008), using the public domain image analysis program ImageJ 1.52i (https://imagej.nih.gov/ij/). For measurement of cell death in roots, seedlings were stained with 10 µg/mL PI (Sigma-Aldrich) for 5 min before imaging.

### Size-exclusion chromatography

Gel filtration experiments were performed using an ÄKTA Fast Protein Liquid Chromatography (FPLC) system (Amersham Biosciences) at 4°C. The absorbance was monitored at 280 nm. The column was calibrated using the molecular weigth standard (29-669 kDa). For recombinant GRXS17 (wild-type and mutated forms), a Superose 6 column was used, equilibrated with a 50 mM Tris-HCl (pH 7.5) buffer containing 150 mM NaCl and 5 mM MgCl_2_ at a flow rate of 0.5 ml min^−1^. All recombinant protein samples were dialyzed before application. For reducing conditions, samples were incubated with 100 mM of DTT and dialyzed against the above-described buffer supplemented with 5 mM of DTT to maintain the proteins in their reduced state.

For separation of total proteins extracts, a HiPrep column Sephacryl S-300 HR (GE Healthcare Life Sciences) was used, equilibrated with a 50 mM Tris-HCl (pH 7.5) buffer containing 150 mM NaCl, 5 mM MgCl_2_, 10% (v/v) glycerol and 0.1% NP40 at a flow rate of 0.5 ml min^−1^. Samples were extracted in the same buffer containing 10 µM of MG132 and anti-protease cocktail (Roche) at a concentration of 1 tablet for 10 mL. Extraction is realized with a ratio of 1 g of plant powder in two volumes of buffer.

### Protein separation by gel electrophoresis and immunoblotting

Precast SDS-PAGE or native gels were used according to the manufacturer’s instructions (Biorad). For immunoblot analysis, gels were transferred to nitrocellulose membranes. Rabbit polyclonal antibodies against GRXS17 (1:25,000) were used for protein Western blotting. Goat anti-rabbit antibodies conjugated to horseradish peroxidase (1:10,000) were used as secondary antibodies and revealed with enhanced chemiluminescence reagents (Immobilon Western Chemiluminescent HRP Substrate, Millipore).

### Thermotolerance assays

For the thermotolerance assays, we applied three different protocols: LAT (Long-Term Acquired Thermotolerance), SAT (Short-Term Acquired Thermotolerance) and TMHT (Thermotolerance to Moderately High Temperatures) as defined by (Yeh et al., 2012). The seeds germinated and grown in horizontal plates containing medium MS (4.4 g/L Murashige and Skoog, 1% (w/v) sucrose, pH 5.8). The percentage of viability was calculated by counting the plants that recovered leaf growth after heat stress. For examining the impact of heat stress on roots, seeds were grown on vertical plates and root growth was daily measured using ImageJ.

### Immunoprecipitation

Proteins from plants expressing the *Pr35S:GRXS17-Flag/HA* or the *Pr35S:GRXS17* constructs were used for immunoprecipitation experiments. The seeds were germinated and grown in vertical plates containing medium ½ MS during 14 days at 20°C and treated or not for 2h at 35°C. The proteins were extracted and submitted to immunoprecipitation. Agarose beads with covalently attached anti-Flag antibody were used according to manufacturer’s instructions and bound proteins were eluted with 0.1 M glycine pH 2.5. In another set of experiments, proteins extracts were first submitted to SEC as described previously. Fractions 18-21 corresponding to oligomers for plants grown at 20°C and fractions 16-19 for plants treated 2 h at 35°C (see Figure 2) were collected and subjected to immunoprecipitation.

### Quantitative Mass Spectrometry Analyses

#### Chemicals and enzymes

Proteomics grade Trypsin and ProteaseMax surfactant were purchased from Promega (Charbonnieres, France). Reversed-phase C18 spin columns, precolumns, and analytical columns were all obtained from Thermo Scientific (Les Ulis, France). Solvents and ion-pairing agents were certified LC-MS grade and all other chemicals were purchased from Sigma-Aldrich (Saint-Quentin Fallavier, France) with the highest purity available.

#### Sample preparation

Immunoprecipitated samples were incubated in the presence of 0.02% (v/v) ProteaseMax with 5 mM DTT for 20 min at 40°C. Free cysteines were further carbamidomethylated by adding 11 mM iodoacetamide for 30 min at room temperature in darkness. Proteins were digested overnight at 35°C with modified porcine trypsin in a 1:50 (w/w) enzyme:substrate ratio. The digestion was stopped by the addition of 0.1% trifluoroacetic acid and peptide mixtures were centrifuged for 10 min at 21,500*xg* at 4°C. Tryptic peptides present in supernatants were then subjected to desalting using reversed-phase C18 spin columns as recommended by the supplier. After elution, desalted peptides were concentrated with a SpeedVac (Eppendorf) and prepared in 3% acetonitrile, 0.1% formic acid (solvent A) were added to obtain a final protein concentration of 0.27 µg/µL based on the initial concentration.

#### Mass spectrometry

Peptide mixtures were analyzed on a Q-Exactive Plus (Thermo Fisher Scientific, San José, CA, USA) coupled to a Proxeon Easy nLC 1000 reversed-phase chromatography system (Thermo Fisher Scientific, San José, CA, USA) using a binary solvent system consisting of solvent A and solvent B (0.1% FA in ACN). 3 µL (800 ng) of tryptic digests were loaded on an Acclaim Pepmap C18 precolumn (2 cm x 75 μm i.d., 2 μm, 100 A) equilibrated in solvent A and peptides were separated on an Acclaim Pepmap C18 analytical column (25 cm x 75 μm i.d., 2 μm, 100 A) at a constant flow rate of 250 nL/min by two successive linear gradients of solvent B from 0% to 25% in 100 min, from 25% to 40% in 20 min and then up to 85% in 2 min followed by an isocratic step at 85% for 7 min. The instrument was operated in positive and data-dependent acquisition modes with survey scans acquired at a resolution of 70,000 (at m/z 200 Da) with a mass range of m/z 375-1,400. After each full-scan MS, up to 10 of the most intense precursor ions (except +1, +5 to +8 and unassigned charge state ions) were fragmented in the HCD cell (normalized collision energy fixed at 27) and then dynamically excluded for 60 s. AGC target was fixed to 3×10^6^ ions in MS and 10^5^ ions in MS/MS with a maximum ion accumulation time set to 100 ms for MS and MS/MS acquisitions. All other parameters were set as follows: capillary temperature, 250°C; S-lens RF level, 60; isolation window, 2 Da. Acquisitions were performed with Excalibur software (Thermo Fisher Scientific, San José, CA, USA) and to improve the mass accuracy of full-scan MS spectra, a permanent recalibration of the instrument was allowed using polycyclodimethylsiloxane ((C_2_H_6_SiO)_6_, m/z 445.12003 Da) as lock mass.

#### Proteomic data processing

MS raw data of HMW complexes were processed with ProteomeDiscoverer 2.2 (Thermo Fisher Scientific, San José, CA, USA) and searched against the UniprotKB *A. thaliana* database (25/10/2015; 31480 entries) combined with a database of classical contaminants using an in-house Mascot search server (Matrix Science, London, UK; version 2.4). Mass tolerance was set to 10 ppm for the parent ion mass and 20 mmu for fragments and up to two missed cleavages per tryptic peptide were allowed. Methionine oxidation, deamidation of asparagine and N-terminal acetylation were taken into account as variable modifications and cysteine carbamidomethylation as a fixed modification. Peptide and protein False Discovery Rates (FDRs) were determined by searching against a reversed decoy database. Peptide identifications were filtered at 1% FDR using the Percolator node. Proteins were filtered at 1% FDR and reverse and contaminants proteins were removed.

MS raw data were processed with MaxQuant software (version 1.6.0.13) and with the associated Andromeda search engine using the UniprotKB *Arabidopsis thaliana* database (25/10/2015; 31480 entries) combined with the MaxQuant database of common contaminants. First and main searches were performed with precursor mass tolerances of 20 and 4.5 ppm, respectively. The minimum peptide length was set to seven amino acids and specificity for protein digestion was restricted to trypsin/P cleavage with a maximum of two missed cleavages per peptides. Methionine oxidation and N-terminal acetylation were specified as variable modifications, carbamidomethylation of cysteines was specified as a fixed modification. The peptide and protein false discovery rate (FDR) was set to 1%. To increase protein identification, a match between runs was systematically performed (Match time window 0.7 min; alignment time window: 20 min) and for label-free quantification of immuno-precipitated proteins, the LFQ algorithm and normalization were chosen and unique and razor peptides were used.

MaxQuant output was further processed and analyzed using Perseus software (version 1.6.07). Protein quantification was performed using the ProteinGroups.txt file after removing reverse and contaminant proteins and the Log2 transformation of quantitative data. Proteins showing at least two unique and razor peptides and having quantitative data in at least three biological replicates were considered for statistical analysis of each temperature condition using a Benjamini-Hochberg-corrected Student T-test. Proteins showing a FDR < 0.05 were considered significantly enriched. For both temperature conditions, proteins having quantitative data in biological samples expressing the tagged GRXS17 but not in the five corresponding control samples were treated separately. In this case, only proteins showing quantitative data in three out of the five biological replicates with at least two unique and razor peptides from samples expressing tagged GRXS17 were considered as significant.

### Circular Dichroism

In preparation for CD analysis, samples of GRXS17 were buffer-exchanged into 20 mM sodium phosphate, 100 mM NaF, pH 7.3, using Bio-Spin^®^ 6 columns, adjusted to 4 μM, and treated with 400 μM of either TCEP or H_2_O_2_ for 1 h at 25°C. Samples were centrifuged and vacuum-degassed for 10 min prior to analysis by CD. CD spectra were collected at 20°C using a J-715 spectropolarimeter (JASCO) with a 0.1 mm path-length cuvette, from a range of 250-185 nm in 1.0 nm sampling and spectrum averaged over 8 repeats. The spectra of the buffer alone with TCEP/H_2_O_2_ was subtracted prior to the conversion of data to units of mean residue ellipticity. Secondary structure content was estimated using the online server, BeStSel (Micsonai et al., 2015). For monitoring change in secondary structure over time, CD spectra of 250-190 nm were collected at 100 nm min^-1^. The secondary structure change of GRXS17 following the addition of a 100-fold excess of H_2_O_2_ was measured as the change in [θ] at 222 nm over a course of 65 min, spectra collected every 41 s. In Prism (GraphPad) a one-phase decay model was fitted to these data, and a rate of conformational change was determined.

## Supporting information

MS/MS analyses

## AUTHOR CONTRIBUTION

LM, JK, JSV, SDL, RL, JM, RS, JPR and CR designed research; LM, JK, LB, AD, SAF, CHM, DY, NHTD, and AD performed research; SAF, CHM, DY, NHTD, SDL, RL and JM contributed new reagents/analytic tools; LM, JK, AD, SAF, CHM, DY, JSV, SDL, RL, JM, RS, JPR, and CR analyzed data; and LM, JK, SAF, DY, JSV, SDL, RL, JM, RS, JPR and CR wrote the paper.

## ACKNOWLEDGMENTS

The authors would like to thank Marion Hamon for technical assistance for mass spectrometry analyses, and Dr. Yee-yung Charng (Academia Sinica, Taiwan) for pioneer experiments on heat stress regimes. This work was supported in part by the Centre National de la Recherche Scientifique, by the Agence Nationale de la Recherche (ANR-Cynthiol 12-BSV6-0011 and ANR-REPHARE 19-CE12-0027), by LABEX DYNAMO ANR-LABX-011 and EQUIPEX CACSICE ANR-11-EQPX-0008, notably through funding of the Proteomic Platform of IBPC (PPI). Laura Martins and Avilien Dard are supported by a Ph.D. grant from the Université de Perpignan Via Domitia (Ecole Doctorale Energie et Environnement ED305). Work performed in the lab of Joris Messens was supported by the Research Foundation-Flanders Excellence of Science project no. 30829584, the Research Foundation-Flanders grants no. G0D7914N, and the VUB Strategic Research Programme (SRP34). We gratefully acknowledge the Core Facilities of Protein Spectroscopy and Protein Biochemistry of Philipps-Universität Marburg. Work performed in the lab of Renate Scheibe was supported with funds from the DFG (SCHE 217; SPP 1710).

## SUPPORTING INFORMATIONS

**Supplemental Figure 1:**
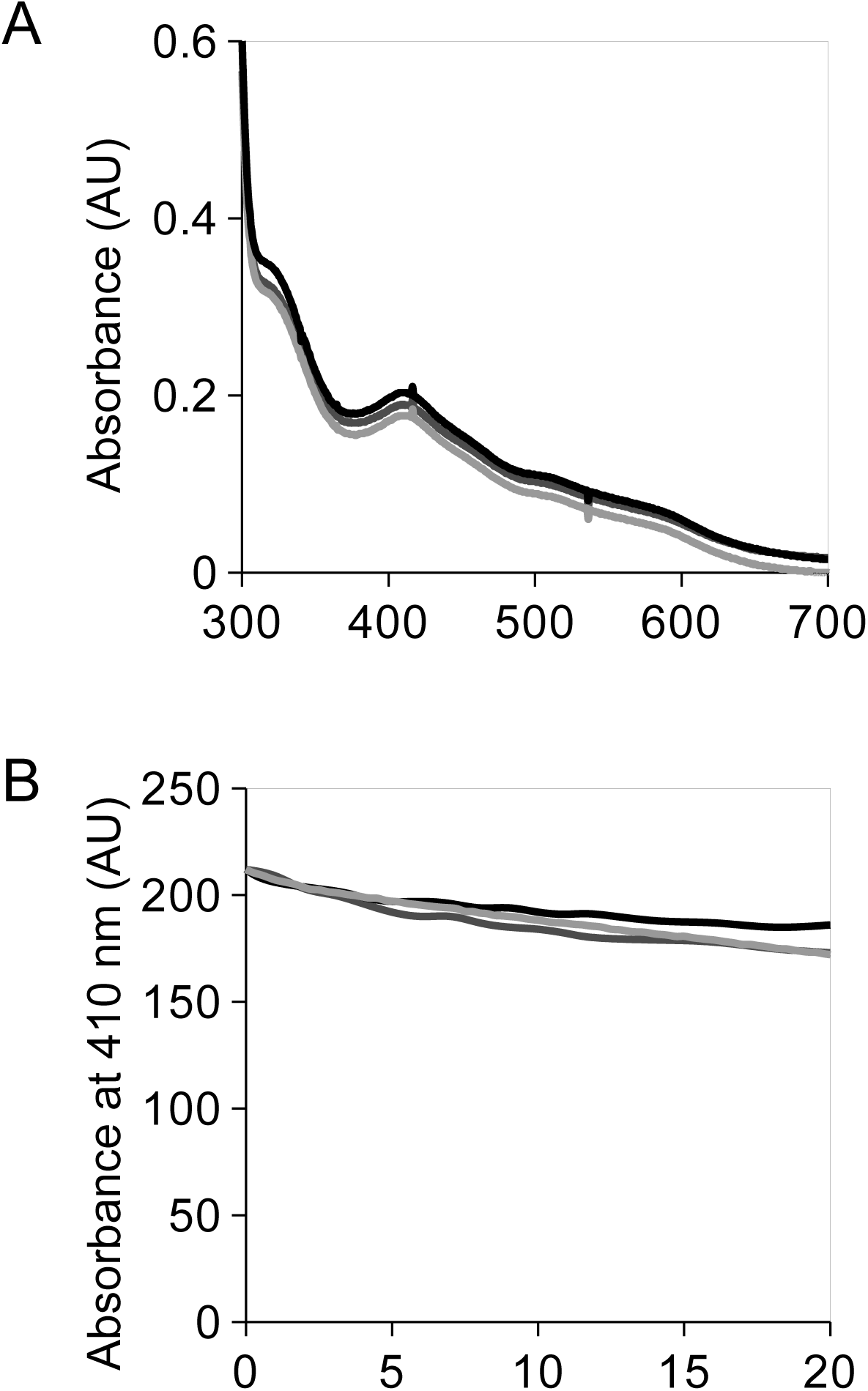
Fe-S cluster stability under increasing temperature. **(A)** Absorption spectra of reconstituted holo-GRXS17 at 25°C (black) or subjected to 35°C (dark grey) or 43°C (light grey). **(B)** Relative absorption at 410 nm of GRXS17 subjected to the same treatments as in **(A).** Each experiment is representative of three biological repetitions.

**Supplemental Figure 2:**
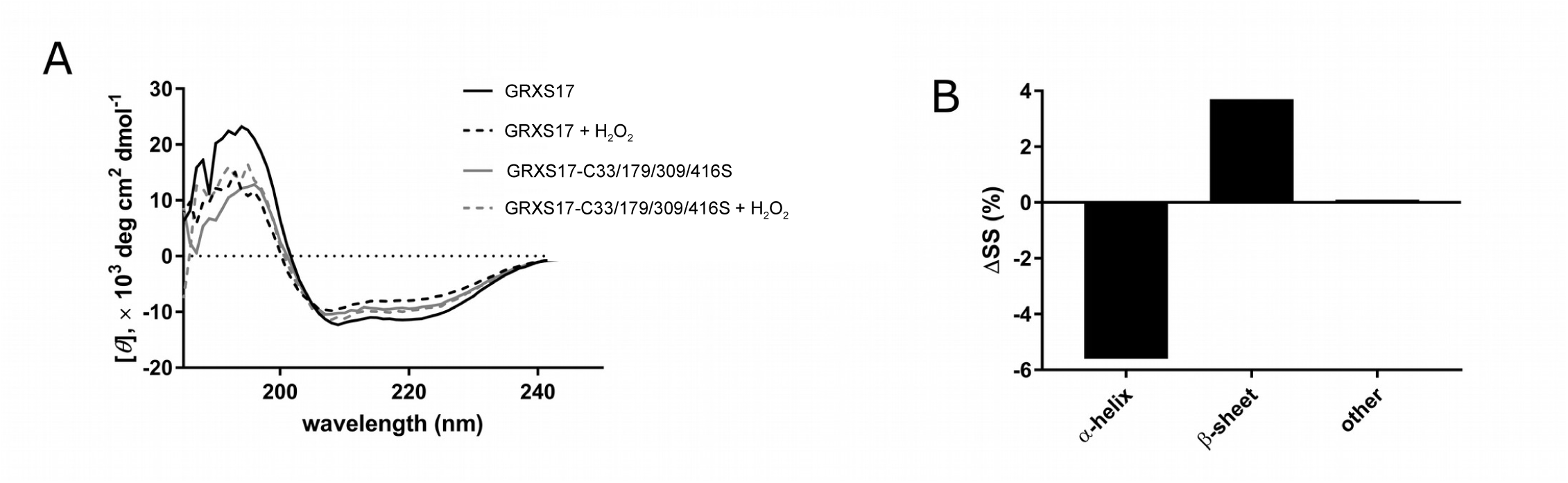
CD study of redox-dependent structural changes in GRXS17 and GRXS17-C33/179/309/416S. (**A**) H_2_O_2_ treatment induces a change in the CD-spectrum of wildtype (GRXS17), but not GRXS17-C33/179/309/416S relative to TCEP-treated sample. (**B**) Estimation of secondary structure content from the obtained CD spectra was performed in BeStSel (Micsonai et al., 2015), and the difference in structural content between the reduced and oxidized forms of GRXS17 determined. Panel B plots the subtraction of the TCEP-treated GRXS17 secondary structure composition from that of the H_2_O_2_-treated sample, thereby indicating the relative change in secondary structure content upon oxidation. Samples of GRXS17 were treated with TCEP or H_2_O_2_ at 400 µM (molar ratio of 100:1 compound to protein).

**Supplemental Figure 3:**
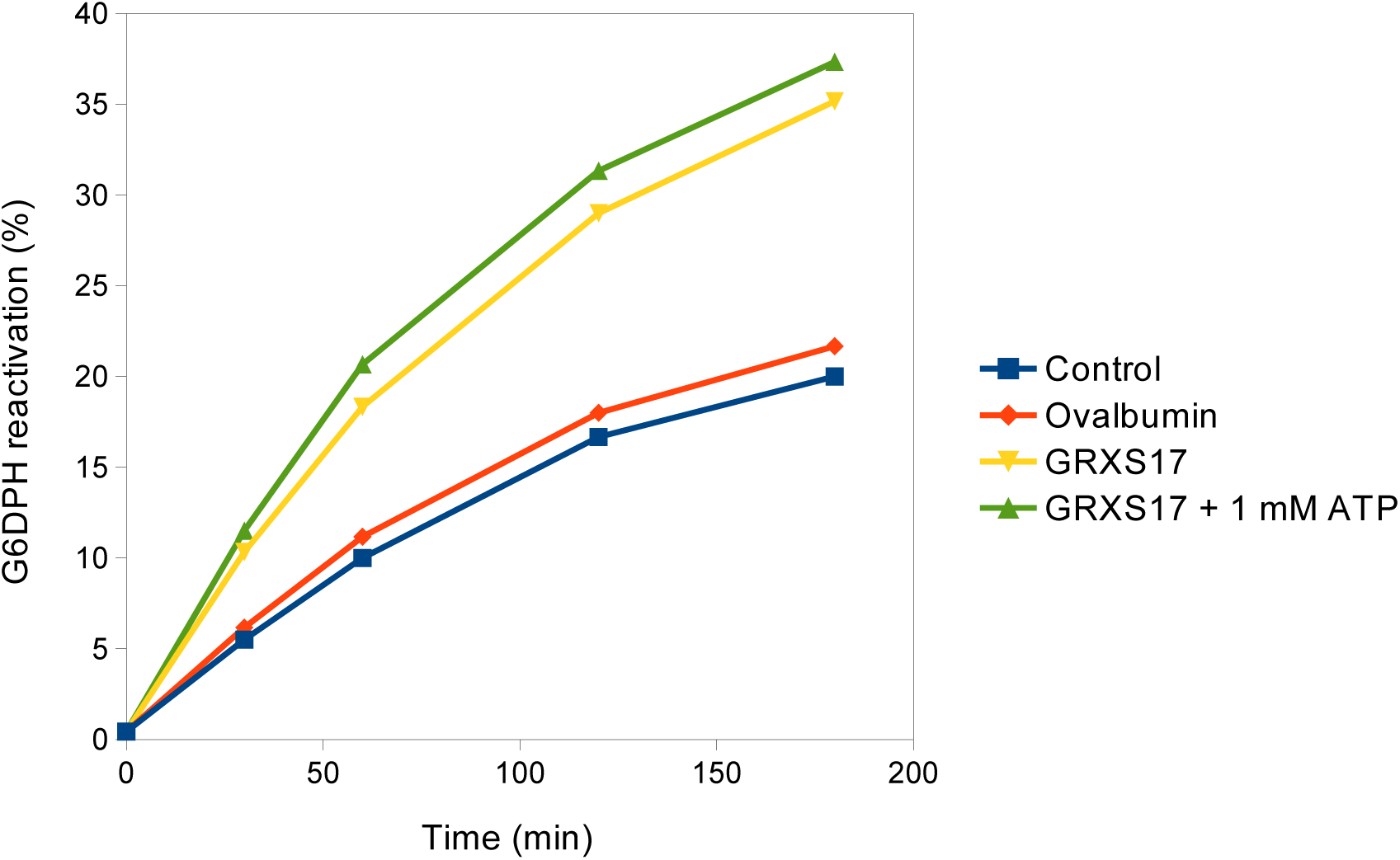
GRXS17 foldase chaperone activity is independent of ATP. GRXS17 foldase activity was measured by reactivation of denaturated glucose-6 phosphate dehydrogenase (G6PDH) at a 5:1 ratio at 30°C. GRXS17 (yellow), but not ovalbumin (red) reactivates G6PDH (dark blue). Addition of ATP (1 mM) to GRXS17 does not significantly modify the foldase activity (green). Data are representative of three independent experiments.

**Supplemental Figure 4:**
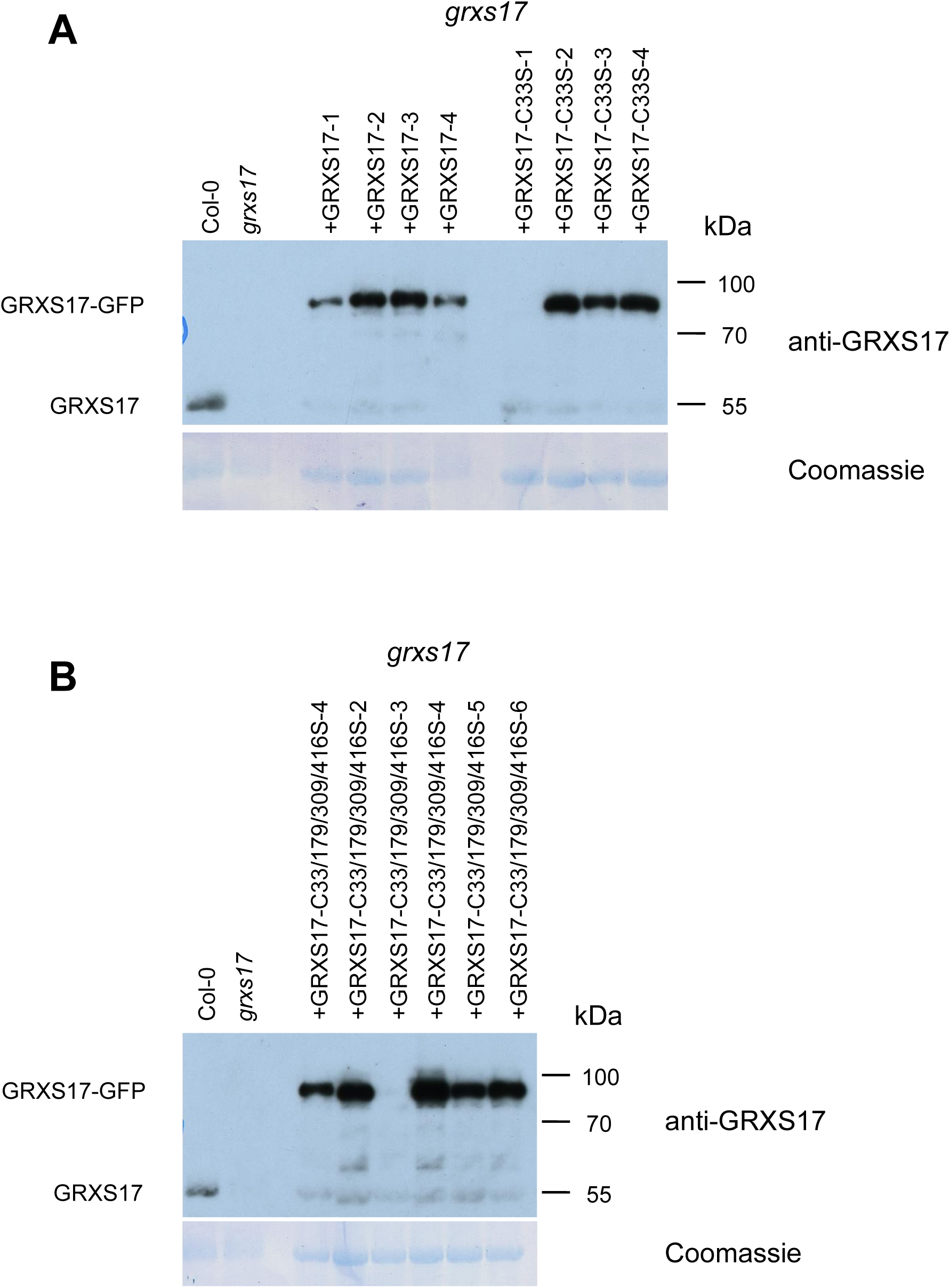
GRXS17 protein expression in complemented grxs17 lines. Total protein extracts were prepared from 2 weeks old plantlets. Samples were prepared from wild-type (Col-0), grxs17 homozygous mutant, and independant grxs17 mutant lines complemented with a (A) Pr35S:GRXS17:GFP construct (+GRXS17), Pr35S:GRXS17-C33S:GFP construct (+GRXS17-C33S) and (B) Pr35S:GRXS17-C33/179/309/416S:GFP construct. Protein extracts were separated by SDS-PAGE and probed with antibodies directed against GRXS17. In Col-0, the 50 kDa signal corresponds to the native GRXS17 protein. In the complemented lines, the 80 kDa signal corresponds to the GRXS17:GFP fusion protein.

**Supplemental Figure 5:**
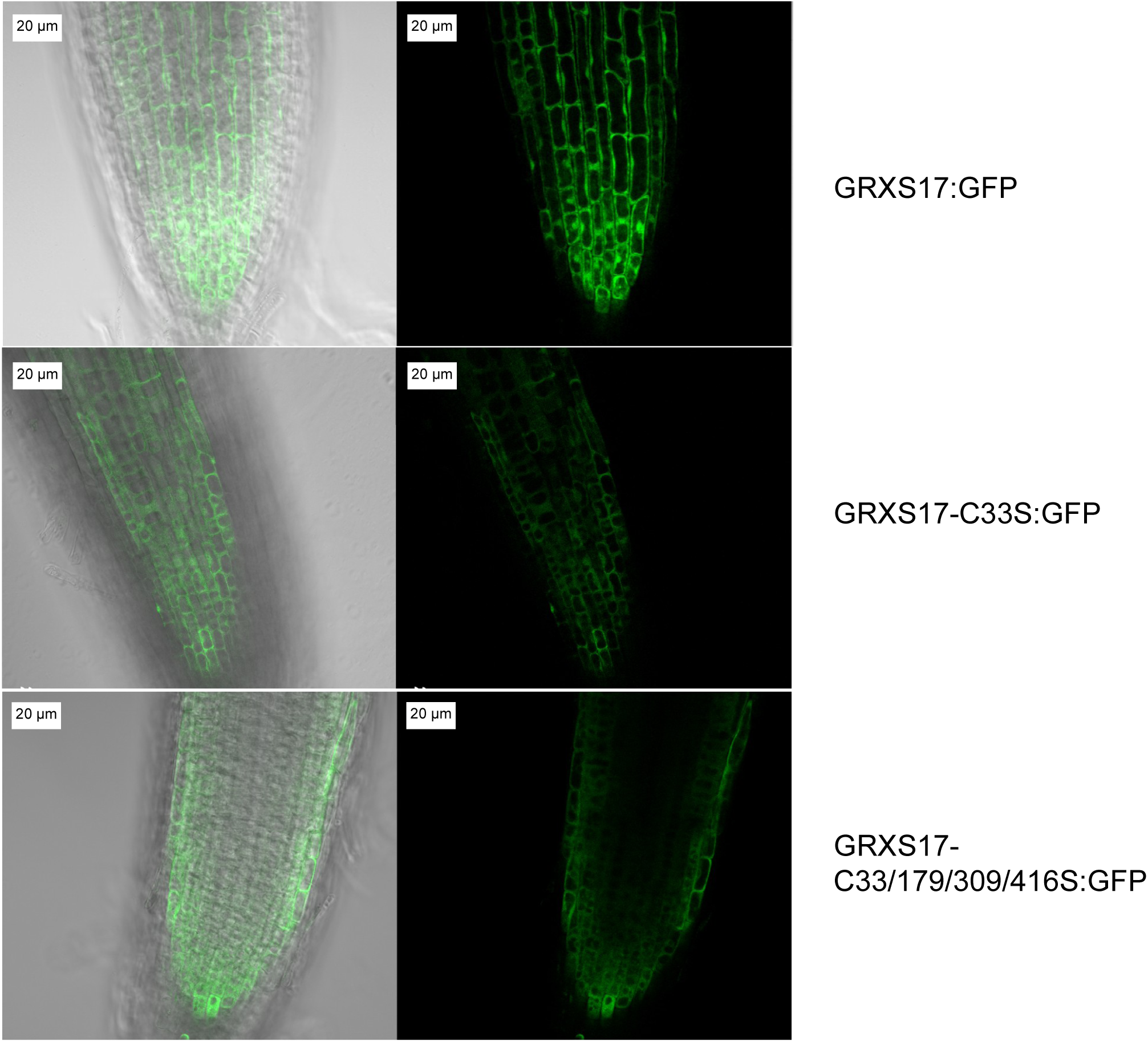
GRXS17:GFP protein expression in complemented *grxs17* lines. GRXS17:GFP fluorescence was monitored by confocal microscopy in root tips from 2 weeks old plantlets. GRXS17:GFP, GRXS17-C33S:GFP and GRXS17-C33/179/309/416S:GFP represent *grxs17* mutant lines which were complemented with GRXS17:GFP, GRXS17-C33S:GFP and GRXS17-C33S/C179S/C309S/C416S:GFP fusion proteins, respectively. Similar expression patterns were observed with other complemented lines.

**Supplemental Figure 6:**
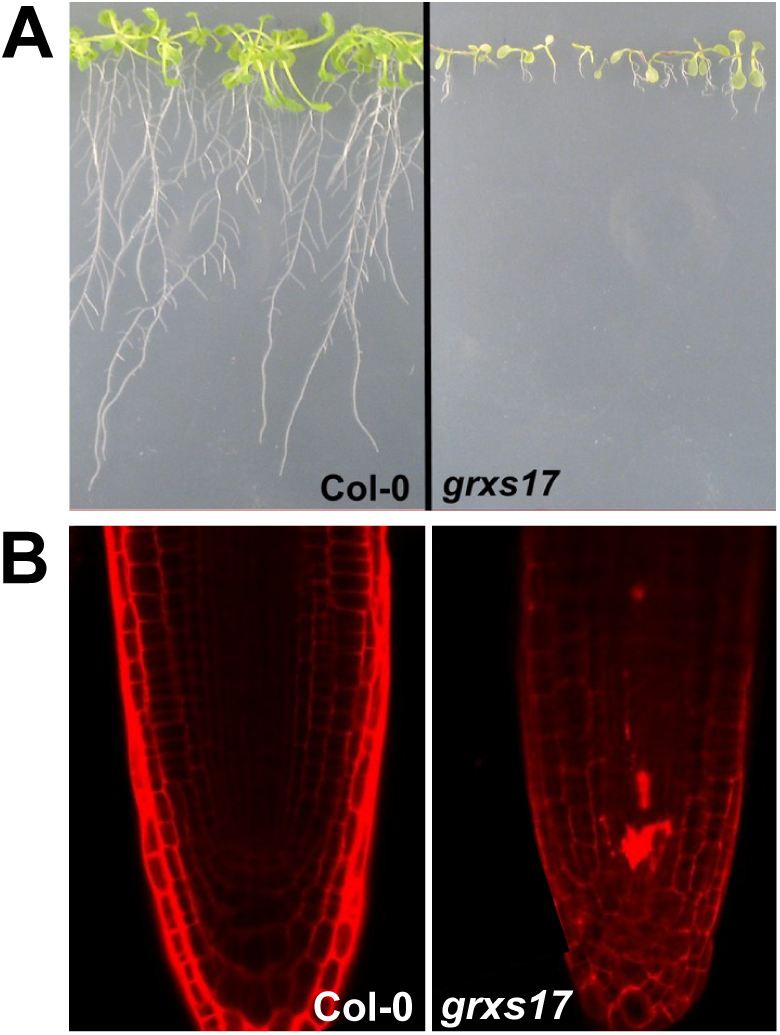
Cells death in the root apical meristem of *grxs17*. Ten day old wild type (Col-0) and *grxs17* mutant plants grown continuously under 28°C. **(A)** Root development. **(B)** Detection of cell death by propidium iodide staining after 10 days of growth.

**Supplemental Figure 7:**
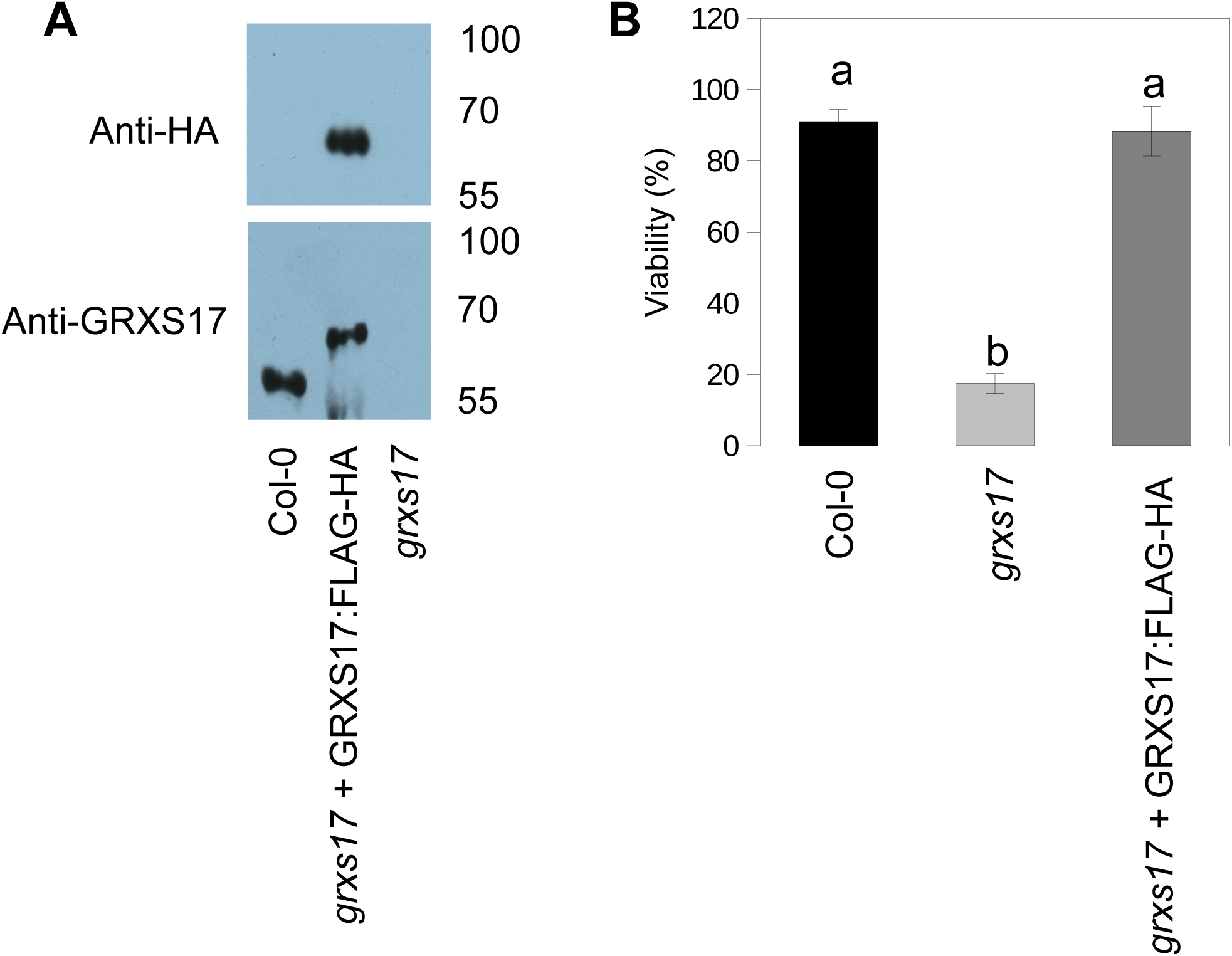
GRXS17:FLAG-HA protein accumulation and TMHT response in *grxs17* complemented lines. (**A**) Total protein extracts were prepared from 2 weeks old plantlets. Samples were prepared from wild-type (Col-0), *grxs17* homozygotous mutant, and *grxs17* complemented with a *Pr35S:GRXS17:FLAG-HA* construct. Protein extracts were separated by SDS-PAGE and probed with antibodies directed against HA or GRXS17 respectively. In Col-0, the ∼55 kDa signal corresponds to the native GRXS17 protein. In the complemented lines, the ∼60 kDa signal corresponds to the GRXS17:FLAG-HA fusion protein. (**B**) TMHT complementation of the *grxs17* mutant by the GRXS17:FLAG-HA protein. Data are means of 4-10 biological repetitions +/- SE, n = 20-25. The letters a and b indicate a significance of p>0.1 and p<0.00001 respectively by the Student’s t-test.

**Supplemental Figure 8:**
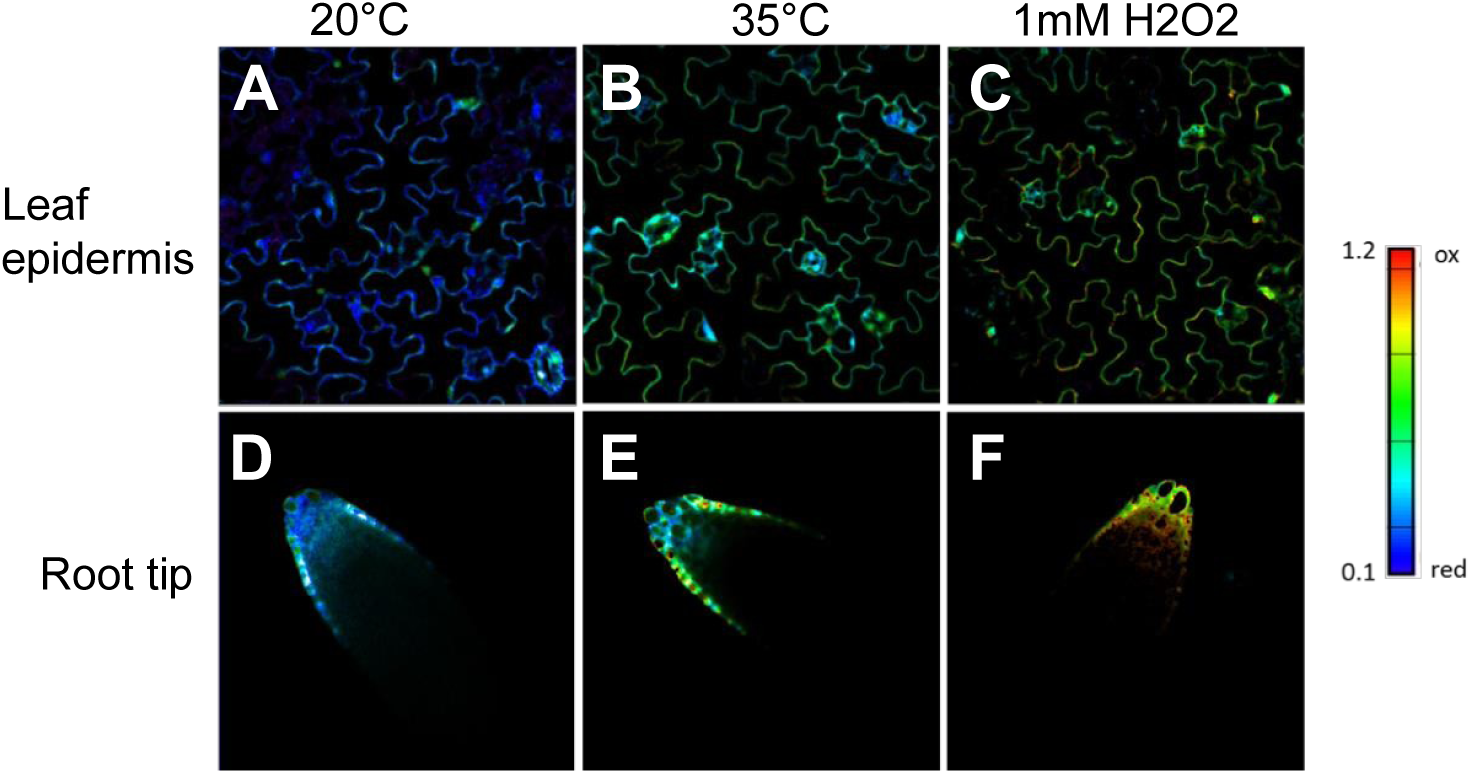
Oxidation of glutathione in the cytosol and nucleus upon high temperature. Images of wild-type leaves (A-C) and root tips (D-F) expressing GRX1-roGFP2 in the cytosol and nucleus were taken by CLSM with excitation at 405 and 488 nm, respectively. GRX1-roGFP2 showed a significant increase in the 405/488nm fluorescence ratio after 1 h treatment at 35°C, as compared to nonstress conditions (20°C) where cytosolic and nuclear glutathione is present mainly in its reduced form. As expected, treatment with 1 mM H_2_O_2_ led to more pronounced oxidation of the GRX1-roGFP2. Single images were used for the calculation of the ratio images. (A and D) Plant grown on 1/2 MS medium at 20°C (B and E) Same plants after 1h treatment at 35°C. (C-F) Plant grown at 20°C with 1 mM H_2_O_2_.

**Supplemental Table 1:** See Supplemental files

**Supplemental Table 2:**
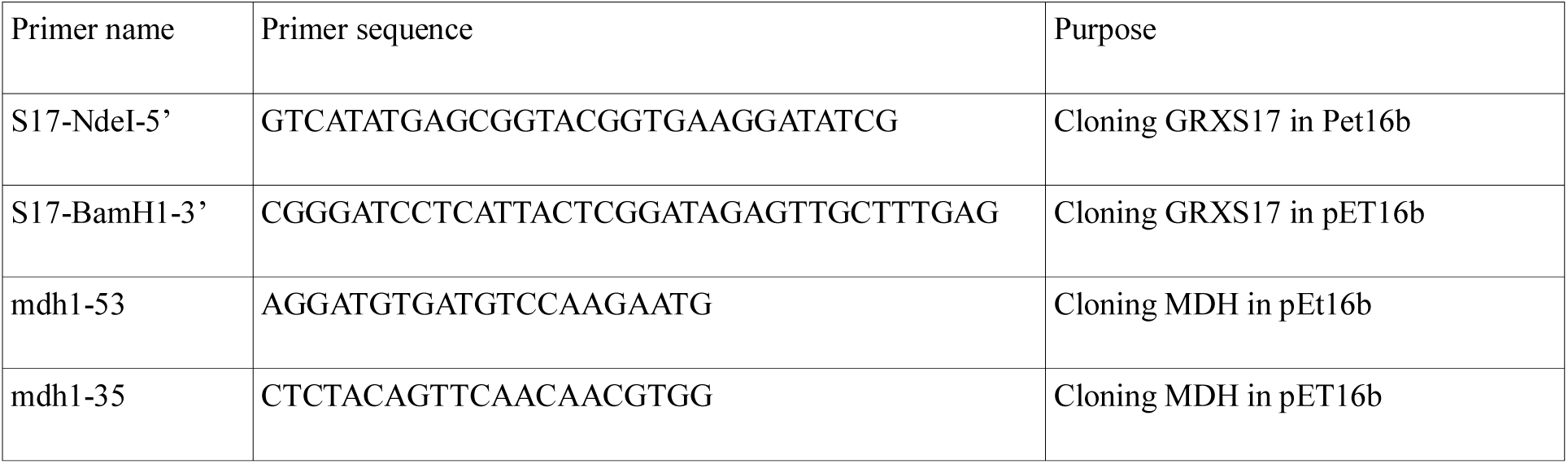
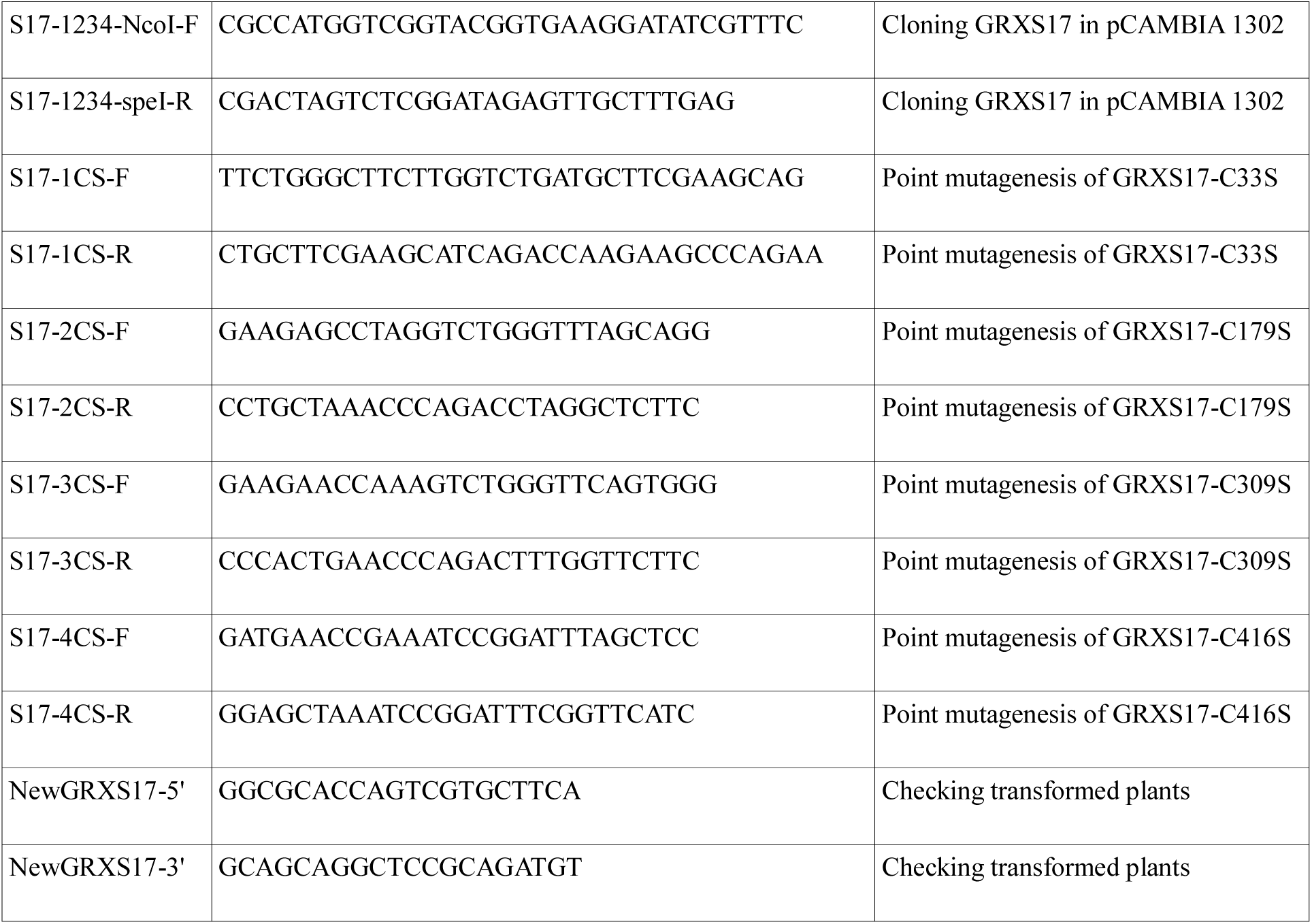
Primers used in this study

